# Intronization enhances expression of S-protein and other transgenes challenged by cryptic splicing

**DOI:** 10.1101/2021.09.15.460454

**Authors:** Kärt Tomberg, Liliana Antunes, YangYang Pan, Jacob Hepkema, Dimitrios A. Garyfallos, Ahmed Mahfouz, Allan Bradley

## Abstract

The natural habitat of SARS-CoV-2 is the cytoplasm of a mammalian cell where it replicates its genome and expresses its proteins. While SARS-CoV-2 genes and hence its codons are presumably well optimized for mammalian protein translation, they have not been sequence optimized for nuclear expression. The cDNA of the Spike protein harbors over a hundred predicted splice sites and produces mostly aberrant mRNA transcripts when expressed in the nucleus. While different codon optimization strategies increase the proportion of full-length mRNA, they do not directly address the underlying splicing issue with commonly detected cryptic splicing events hindering the full expression potential. Similar splicing characteristics were also observed in other transgenes. By inserting multiple short introns throughout different transgenes, significant improvement in expression was achieved, including >7-fold increase for Spike transgene. Provision of a more natural genomic landscape offers a novel way to achieve multi-fold improvement in transgene expression.

## Introduction

Efficient expression of exogenous genes to produce recombinant therapeutic human proteins and immunogens from DNA expression vectors in mammalian cells *in vitro* and *in vivo* is of fundamental importance to the biopharmaceutical industry. Poor expression has been reported for many recombinant proteins and for both SARS-CoV and SARS-CoV-2 Spike (S) proteins when using non-optimized wildtype complementary DNA (cDNA) sequences (1, 2). In many cases expression of exogenous proteins has been iteratively improved by increasing translational efficiency via codon optimization of the protein coding sequence (CDS) and overall mRNA levels can be enhanced by promoter choice and addition of cis-acting modules like 5’UTR introns.

Codon optimization was first applied over 40 years ago, before the cDNA sequence of the desired peptide hormone was even known to the authors (3). It is now routinely used for applications in bioproduction as well as *in vivo* nuclear therapeutic applications (4, 5). Although codon optimization is now a standard tool used in recombinant protein expression systems to increase expression, potential downsides affecting protein structure and function have also been raised (6, 7).

Given that the number of mRNA transcripts per cell will influence protein levels emphasis has been placed on increasing transgene copy number as this can overcome poor expression and quality (full-length) transcripts. Convenience of packaging and delivery by viral vector systems and the ability to achieve higher copy numbers of smaller constructs has led to the almost universal use of cDNAs by the biopharmaceutical industry even though the importance of intronic sequences for eukaryotic gene expression was shown over 40 years ago (8). While 97% of endogenous protein coding genes in humans contain introns and intron/exon boundaries are extremely highly conserved across species, non-intron containing cDNAs are almost invariably used as transgenes. Numerous aspects of gene expression are affected by introns including initial transcription of the gene, rate of transcription, polyadenylation, nuclear export, RNA editing, translational efficiency, and mRNA decay (9, 10).

Examples of 5’UTR intron mediated expression improvements have led to inclusion of 5’UTR intron in many transgenes’ designs (11). Benefits from adding multiple introns into transgenes are however controversial, while some studies describe benefits achieved with constructs containing multiple endogenous introns (12) and heterologous introns (13–15) other studies report no added benefit achieved by more than a single intron (16).

Expressing genetic sequences in a novel context, such as genes from a cytoplasmic RNA virus that never enters the nucleus as a cDNA is inevitably challenged by cryptic splicing, a situation where a cryptic intron is efficiently recognized and excised within the intended CDS. This can lead to no or low expression of the full-length protein as well as production of truncated and out-of-frame protein fragments (17, 18). Cryptic splice sites (SS) that contain a consensus motif sequence in addition to the canonical splice donor (SD) dinucleotide ‘GT’ or the canonical splice acceptor (SA) dinucleotide ‘AG’ within the CDS can be removed by codon optimization. In situations where desired protein expression is observed from a codon-optimized construct, alternative splicing events that lead to aberrant mRNAs, protein fragments and out of frame products may only be discovered by in-depth analysis as recently reported for SARS-CoV-2 S protein sequences used in some COVID-19 vaccines (19, 20).

Here, we report extensive cryptic splicing events within both wildtype and various codon-optimized SARS-CoV-2 S protein sequences as well as other transgenes. While removing canonical cryptic SS alleviates the repetitive splicing outcomes facilitated by the presence of conserved SS, it does not resolve the issue. We show that by inserting multiple introns protein expression of a number of transgenes has been enhanced by up to 9-fold, opening an opportunity for improving transgene expression and protein quality in bioproduction as well as in DNA vaccines.

## Results

### Poor expression of full-length S protein from the non-optimized DNA sequences

SARS-CoV-2 CDS are conventionally codon-optimized for expression as DNA transgenes. Wildtype (P1, P2) and codon-optimized (P13) constructs for S protein were compared with respect to production of protein (Figure 1). Anti-S protein or FLAG-tag antibodies failed to detect expression from either wildtype construct while the codon-optimized version yielded both full length and variety of smaller products (Figure 1B). Surface staining was detected at a low level from P2 while the codon-optimized construct yielded detectable S protein expression in nearly 60% of cells (Figure 1C). Changes to the 5’ and 3’ expression elements around the transgene CDS did not improve wildtype S expression (P3-P9, Sup Figure 1). However, when the wildtype S protein CDS was transfected as a mRNA, protein expression was detected in 9% of cells confirming the coding sequence was functional. Transfection of mRNA from the P13 codon-optimized construct yielded similar numbers of expressing cells compared with the DNA counterpart (59% vs 53%, Figure 1D).

**Figure 1.**
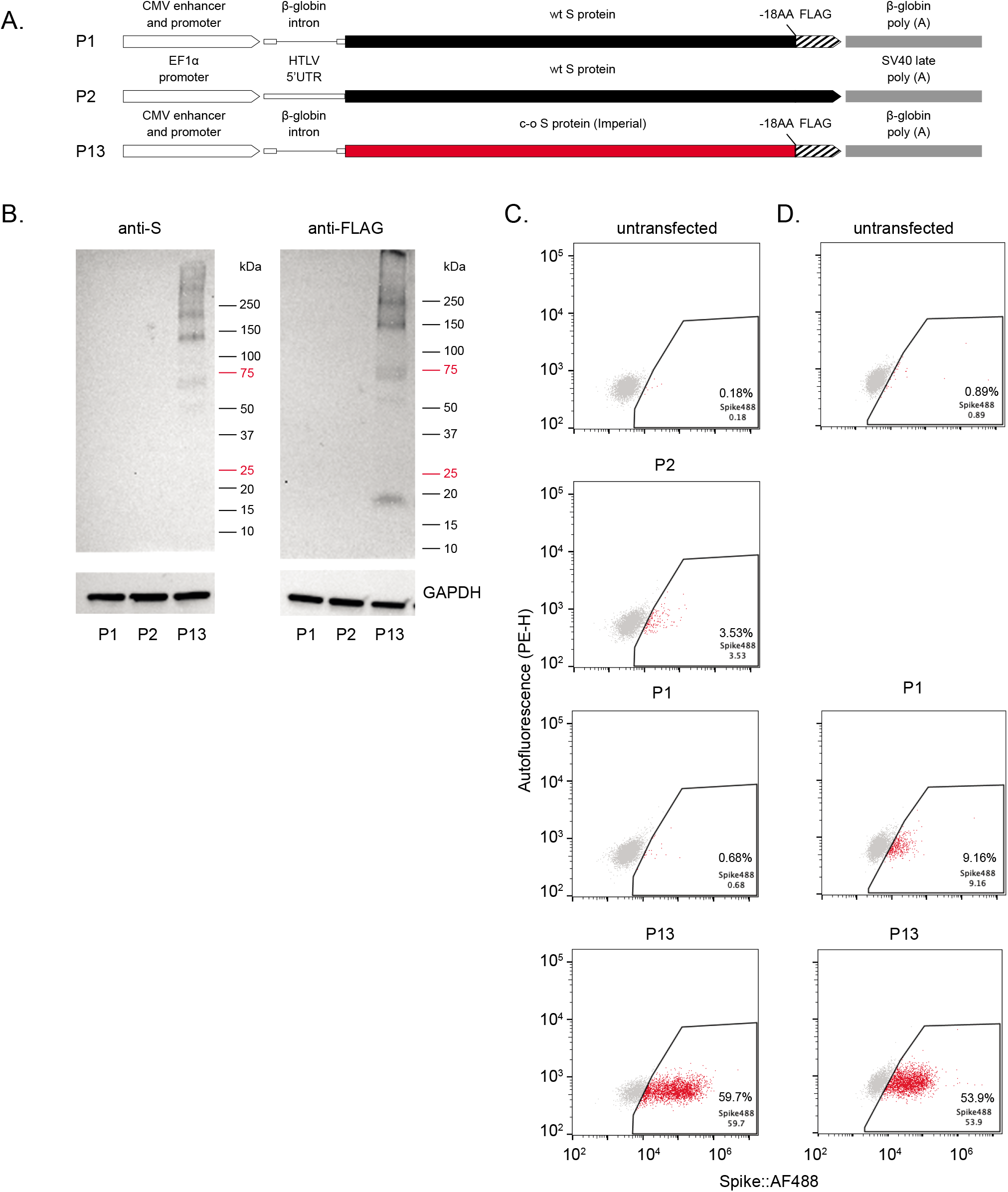
Poor expression of S protein from wildtype (non-optimized) DNA constructs. S protein expression from two different wildtype (wt) S protein constructs P1 (assembled in-house) and P2 (commercially available) was compared with a codon-optimized (c-o) S construct P13 **[A]**. Two Western blots (either anti-S or anti-FLAG antibodies) **[B]** as well as anti-S flow cytometry **[C]** demonstrated detectable expression for the c-o S protein but either undetectible or very low levels of expression for the wt constructs. Expression could be detected when wt S protein sequence was introduced to cells as *in vitro* synthesized mRNA **[D]**. Note: anti-FLAG was not expected to detect P2 construct.

### Non-optimized DNA sequence is cryptically spliced

To understand the discrepancy in expression between wildtype S DNA and mRNA as templates, an RT-PCR spanning from the 5’UTR to 3’UTR was performed on cells transfected with P1. A predominant 280 bp PCR product was detected while the expected 4011 bp product was not observed (Figure 2A, 2C). Sequencing identified this as an alternatively spliced mRNA transcript generated using a SD site upstream of the start codon (SD-15) and a SA site at 3’ end of the S protein CDS (SA3716). A similar outcome, albeit using a different SD site within the Human T-cell Leukemia Virus (HTLV) 5’UTR, was observed for P2 (Sup Figure 2).

**Figure 2.**
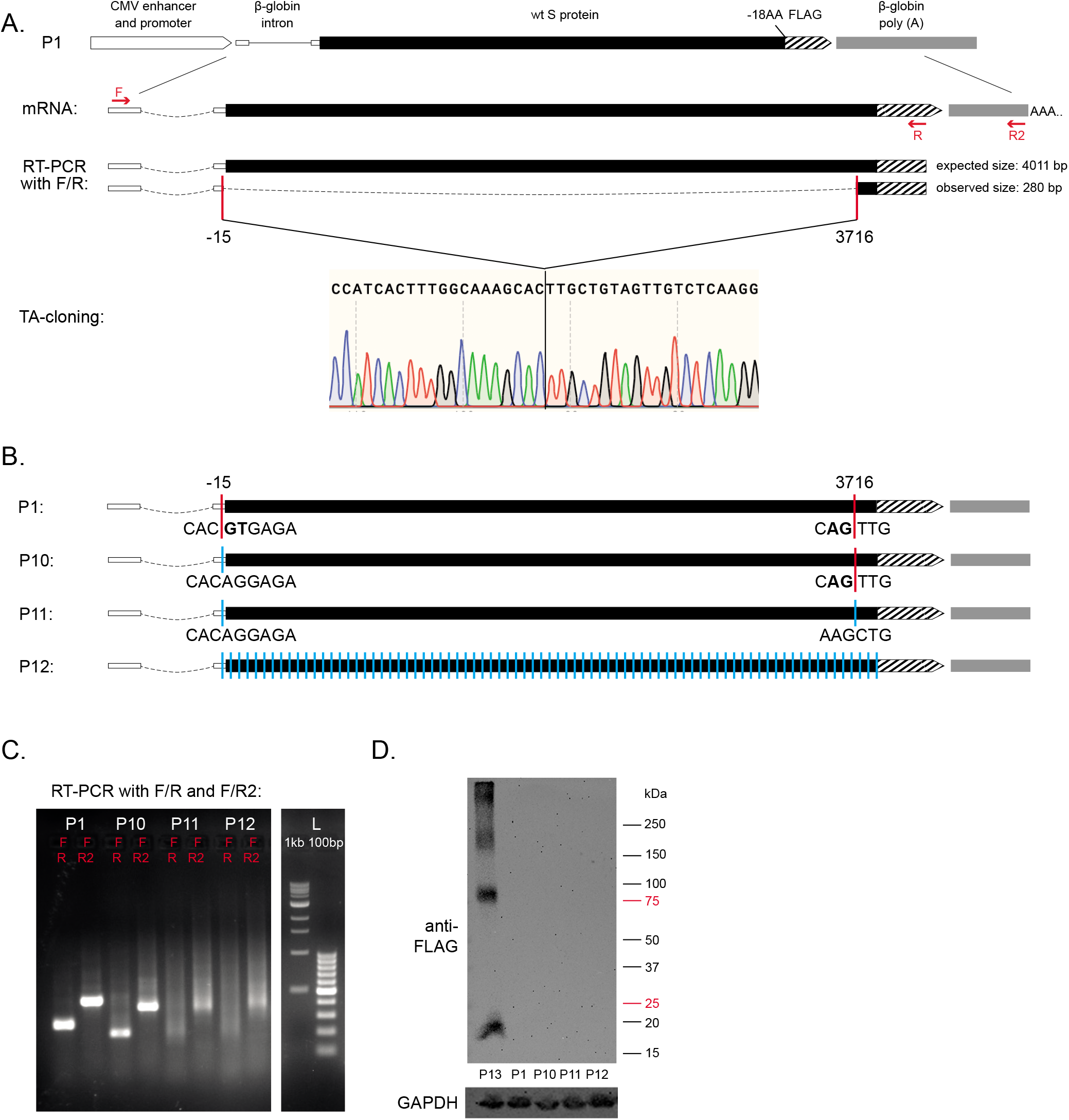
Wt S protein construct leads to cryptic splicing events. RT-PCR with primers positioned from 5’UTR to 3’UTR of the P1 construct (F,R,R2) only highlighted the presence of one preferred alternatively spliced mRNA product (280 bp) instead of the expected full length mRNA (4011 bp). The cryptic SD (−15) and SA (3716) sites for the short transcript were mapped relative to translation start site of S protein using TA-cloning and Sanger sequencing **[A]**. Three further constructs were made to assess the cryptic splicing. First, the cryptic SD (P10) or both SD and SA (P11) sites were removed using codon-optimisation. In addition, a construct with all predicted SS within wt S protein sequence removed (P12) was generated **[B]**. RT-PCR across these new constructs demonstrated no visible full length mRNA product **[C]** and no protein was detected using Western blots **[D]**.

These observations reflect the biology of SARS-CoV-2 whose genome as a cytoplasmic RNA virus has not evolved to support expression from a nuclear DNA template. The virus’s S protein sequence alone harbors 111 SS predicted by ASSP software (Sup Table 1) (21). In an attempt to resolve the cryptic splicing outcome, three constructs eliminating both observed and predicted SS were generated while retaining identical amino acid sequence. P10 has SD-15 site removed, P11 has both SD-15 and SA3716 removed while P12 has 108 out of the 111 predicted SS removed (Sup Table 1, Figure 2B). These edits failed to improve full length S protein expression (Figure 2C-D), but did change the underlying cryptic splicing outcomes from a single preferred event using available canonical SS to a spread of random events stemming from non-canonical SS in P12 (Figure 2C, Sup Figure 3).

**Table 1.** All used DNA constructs

| No | Name | 5' elements | 3' elements |
| --- | --- | --- | --- |
| Wuhan-Hu-1 isolate SARS-CoV-2 S protein sequence |  |  |  |
| P1 | P1_pMD2.G_Wuhan_S18F | CMV promoter + β-globin intron | β-globin polyA |
| P2 | P2_InVivoGen_S | EF-1α promoter +HTLV (Human T-Cell Leukemia Virus) 5'UTR | SV40 polyA |
| P3 | P3_EFS_Wuhan_S18F | EFS promoter | β-globin polyA |
| P4 | P4_EF1a_Wuhan_S18F | EF1α promoter | β-globin polyA |
| P5 | P5_PGK_Wuhan_S18F | PGK promoter | β-globin polyA |
| P6 | P6_CMV_Wuhan_S18F | CMV promoter | β-globin polyA |
| P7 | P7_CMV_Wuhan_S_WPRE | CMV promoter + β-globin intron | WPRE |
| P8 | P8_EFS_Wuhan_S_WPRE | EFS promoter | WPRE |
| P9 | P9_CMV_Wuhan_S_WPRE | CMV promoter | WPRE |
| Splice site optimized SARS-CoV-2 S protein sequences |  |  |  |
| P10 | P10_pMD2.G_Wuhan_S18F_noSD | CMV promoter + β-globin intron | β-globin polyA |
| P11 | P11_pMD2.G_Wuhan_S18F_noSDSA | CMV promoter + β-globin intron | β-globin polyA |
| P12 | P12_pMD2.G_Wuhan_S18F_noSS | CMV promoter + β-globin intron | β-globin polyA |
| Codon-optimized SARS-CoV-2 S protein sequences |  |  |  |
| P13 | P13_pMD2.G_Imperial_S18F | CMV promoter + β-globin intron | β-globin polyA |
| P14 | P14_pMD2.G_Kent_S18F | CMV promoter + β-globin intron | β-globin polyA |
| P15 | P15_pMD2.G_UCSC_S18F | CMV promoter + β-globin intron | β-globin polyA |
| P16 | P16_pMD2.G_NIBSC_S18F | CMV promoter + β-globin intron | β-globin polyA |
| P17 | P17_CMV_Imperial_S | CMV promoter | β-globin polyA |
| P18 | P18_CMV_CanSino_S | CMV promoter | β-globin polyA |
| P19 | P19_CMV_SputnikV_S | CMV promoter | β-globin polyA |
| Codon-optimized MERS-CoV S protein sequence |  |  |  |
| P20 | P20_MERS_Oxford_S | CMV promoter + intron A | β-globin polyA |
| Human ACE2 gene CDS |  |  |  |
| P21 | P21_EF1a_hACE2 | EF1α promoter | β-globin<br>polyA |
| Intronization experiment constructs |  |  |  |
| P22 | P22_CMV_NIBSC_S | CMV promoter | β-globin<br>polyA |
| P23 | P23_CMV_NIBSC_S_3IN | CMV promoter | β-globin<br>polyA |
| P24 | P24_CMV_NIBSC_S_7IN | CMV promoter | β-globin<br>polyA |
| P25 | P25_CMV_NIBSC_S_13IN | CMV promoter | β-globin<br>polyA |
| P26 | P26_CMV_NIBSC_S_14IN | CMV promoter | β-globin<br>polyA |
| P27 | P27_PGK_hACE2 | PGK promoter | bovine GH<br>polyA |
| P28 | P28_PGK_hACE2_3IN | PGK promoter | bovine GH<br>polyA |
| P29 | P29_PGK_hACE2_6IN | PGK promoter | bovine GH<br>polyA |
| P30 | P30_PGK_hACE2_9IN | PGK promoter | bovine GH<br>polyA |
| P31 | P31_PGK_mCherry | PGK promoter | SV40 late<br>polyA |
| P32 | P32_PGK_mCherry_3IN | PGK promoter | SV40 late<br>polyA |
| P33 | P33_PGK_mCherry_7IN | PGK promoter | SV40 late<br>polyA |

**Figure 3.**
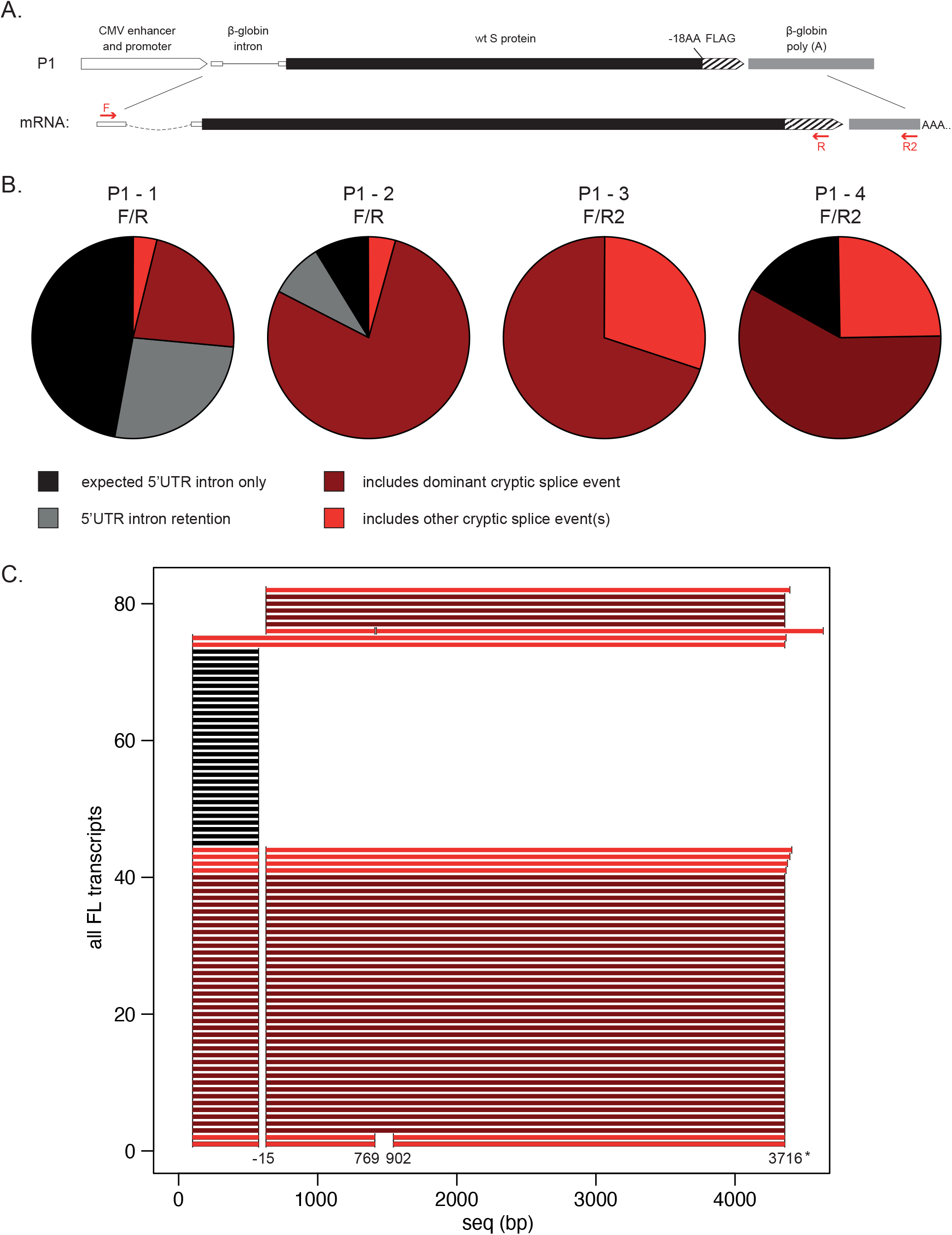
Nanopore cDNA direct sequencing validates dominant splice variant for P1 construct. After cDNA first strand synthesis with either primer R or R2 and second strand synthesis with primer F, four biological replicates were prepared for direct cDNA sequencing **[A]**. Distribution of mRNA splicing outcomes among full-length (FL) reads (≥90% span of the expected sequence) within the replicates **[B]**. Overview of all FL splicing events combined into a single plot. Every event is represented as a single line from its start to end along the sequence length **[C]**. *Co-ordinates relative to translation start site ATG.

### Cryptic splicing validated with Nanopore direct cDNA sequencing

To efficiently detect cryptic splicing events, Nanopore cDNA direct sequencing was carried out on four biological replicates from cells transfected with P1 plasmid and two different transcript specific primers for cDNA synthesis (R and R2, Figure 3A). The distribution of splicing events had some variation. The primer positioned at the most 3’ end of the construct (R2) captured more diverse cryptic splicing compared to one positioned more 5’, as expected (Figure 3B). The most frequent splicing outcome was consistent with the use of SD-15 and SA3716 sites (Figure 3B-C).

### Codon optimization impacts cryptic splicing frequency but does not abolish it

While codon-optimized S transgenes improve S protein expression, this change cannot be solely explained by enhanced protein translation as the wildtype codon sequence in the context of live SARS-CoV-2 viral infection results in ample expression of S protein. Furthermore, as we have shown, removal of canonical SS from the wildtype sequence simply uncovers non-canonical cryptic SS.

Four different codon-optimized S protein sequences P13-P16 with different optimization approaches (Sup Table 2) were assessed to investigate this further (Figure 4A). All constructs resulted in full size S protein expression and surface staining (Figure 4B–4C). Nevertheless, an RT-PCR analysis captured multiple transcripts in addition to the anticipated full-size 4kb product (Figure 4D) and various cryptically spliced events were identified for all constructs (Sup Figure 4). Two types of splicing outcomes were observed: random unique events using various canonical and non-canonical SS (P13, P16) or repetitive splicing outcomes stemming from the same selected canonical SS (P14, P15).

**Table 2.**
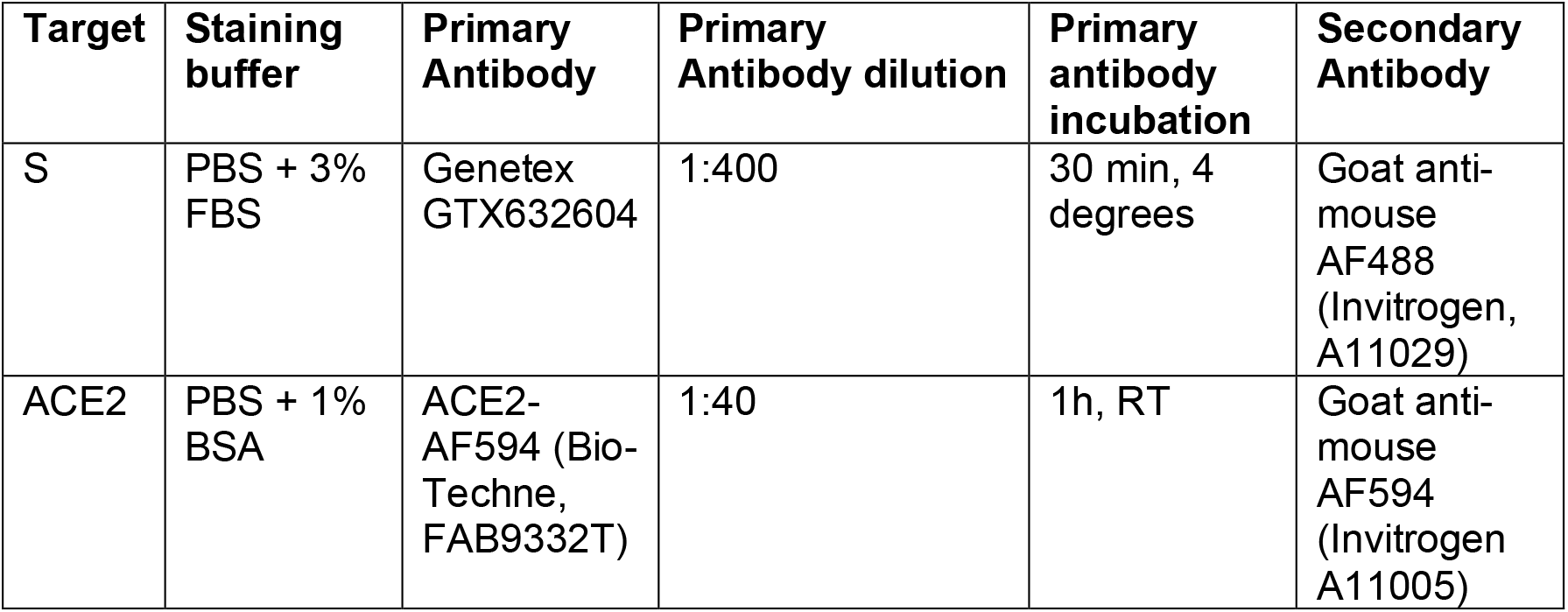
Surface staining antibodies.

| **Target** | **Staining buffer** | **Primary Antibody** | **Primary Antibody dilution** | **Primary antibody incubation** | **Secondary Antibody** |
| --- | --- | --- | --- | --- | --- |
| S | PBS + 3% FBS | Genetex GTX632604 | 1:400 | 30 min, 4 degrees | Goat anti-mouse AF488 (Invitrogen, A11029) |
| ACE2 | PBS + 1% BSA | ACE2-AF594 (Bio-Techne, FAB9332T) | 1:40 | 1h, RT | Goat anti-mouse AF594 (Invitrogen A11005) |

**Figure 4.**
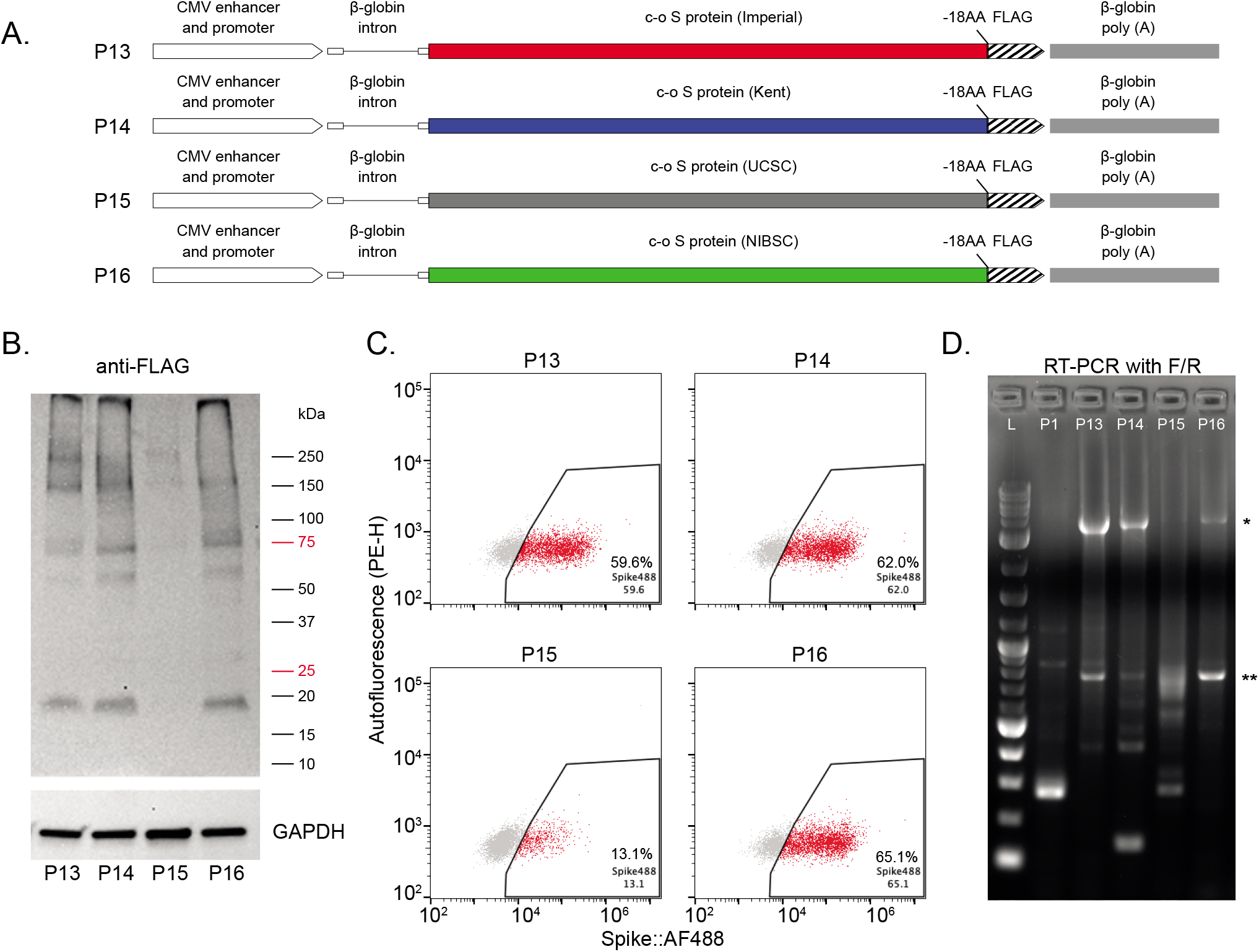
Codon-optimization improves S protein expression with varied success. Four constructs carrying different c-o S protein sequences were generated **[A]**. All c-o constructs had measurable S protein expression observed both with anti-FLAG Western blot **[B]** as well as with anti-S flow cytometry **[C]**. RT-PCR using F/R primers amplified the expected full length mRNA product (*) in most c-o constructs but also some construct-specific alternatively spliced products **[D]**. **mispriming event

Transcripts from P13 and P15 were investigated further by Nanopore cDNA direct sequencing using two different 3’ primers as before (Figure 5A). The overall proportion of correct full-length reads was much higher for P13 (P13-1: 81%, P13-2: 53%) compared to P15 (P15-1: 23%, P15-2: 3%), consistent with the S protein expression levels. Splicing events from P13 were randomly distributed across the sequence with no strong SS preference for SD (orange bars) or SA (grey bars) sites, compared to the SD/SA of the 5’UTR intron (Figure 5C). These SS usage characteristics remained similar also when looking at all the mapped reads as well as when a construct without a 5’UTR intron was used (P17, Sup Figure 5). P15 splicing events repeatedly relied on the same few SS (Sup Table 1) within the sequence observed both in the full length read population (Figure 5D) as well as when all reads were considered (Sup Figure 6A).

**Figure 5.**
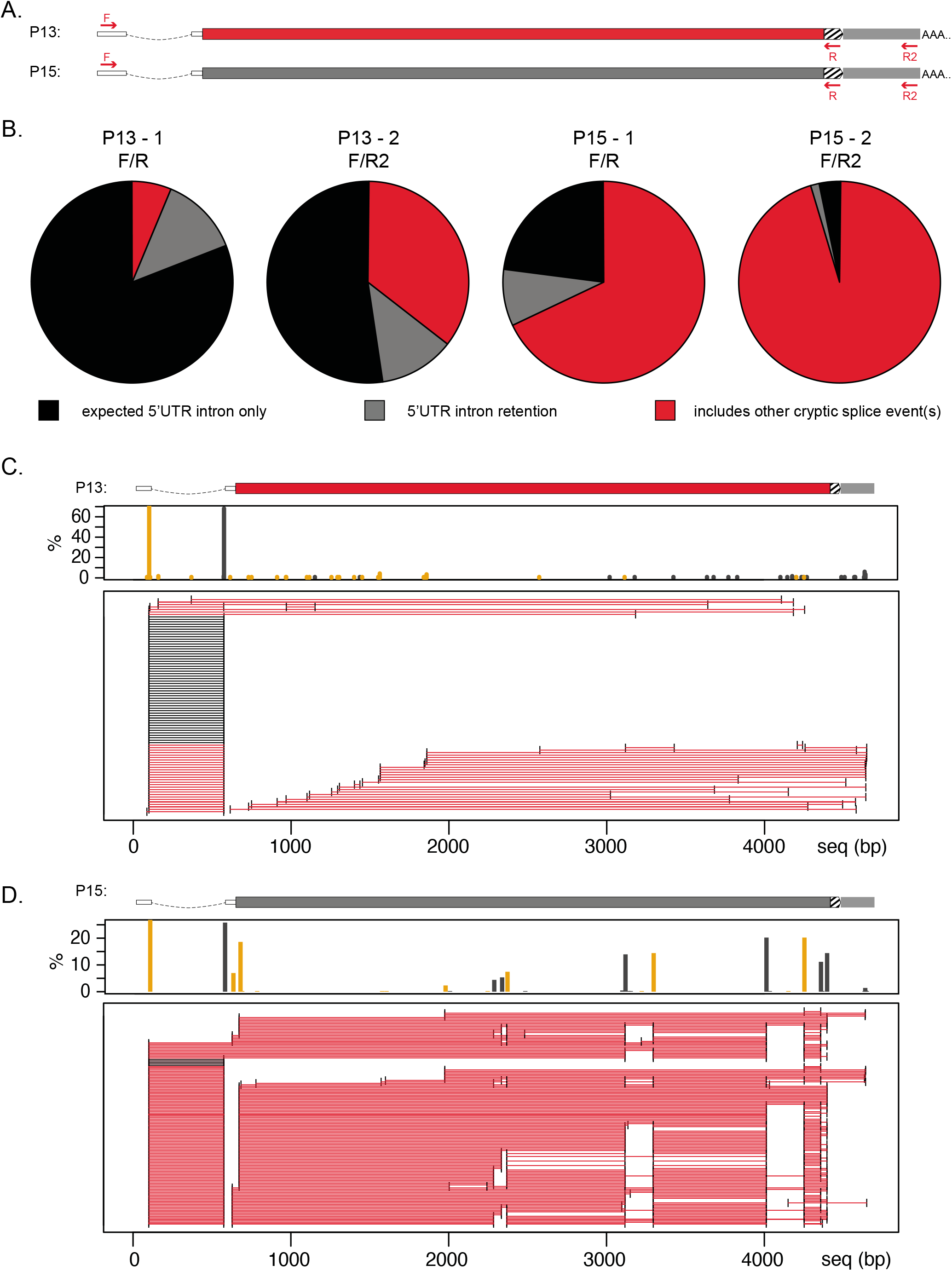
cDNA direct sequencing validates cryptic splicing in c-o S protein constructs. cDNA first strand synthesis with either primer R or R2 and second strand synthesis with primer F carried out for c-o constructs P13 and P15 **[A]**. Distribution of splicing outcomes in FL reads within the c-o samples **[B]**. Overview of all splicing events within FL transcripts and the frequency of using each SD (orange) and SA (grey) site for samples P13-2 **[C]** and P15-2 **[D]**.

**Figure 6.**
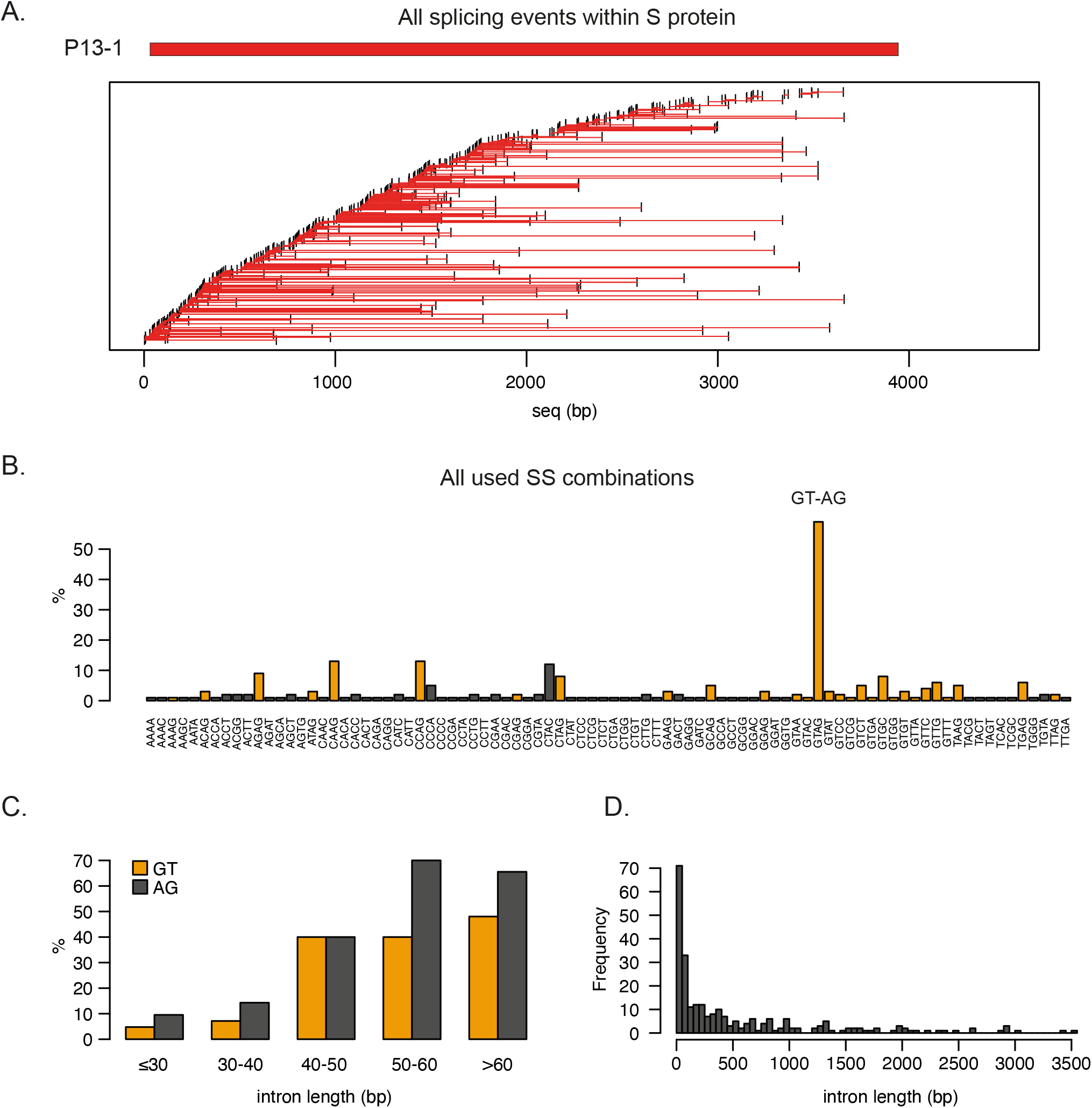
Characteristics of P13 splicing events within S-protein sequence. P13 splicing events are mostly unique and spread across the entire S-protein sequence **[A]**. The majority of these events use canonical SD/SA sites GT-AG or at least one of the two, highlighted in orange **[B]**. Canonical SS-usage increases for introns over 50bp of length **[C]**. Distribution of intron length highlights preference for smaller introns **[D]**.

While both P13 and P15 codon optimization strategies resulted in a similar number of predicted SS within the S protein sequence (78 v 86), the prediction scores across the SS were significantly higher for P15 (t-test: p<0.0005, Sup Table 1). Nevertheless, not all the frequently observed SS in P15 had a high prediction score. Preferred sites included SS originally present in wildtype S sequence, slightly modified wildtype S sequence SS as well as novel sites introduced by the codon optimization process (Sup Figure 6B). Different combinations between the same set of SD/SA sites were observed, but the distribution was not random and often the preferred combination for a SD site was with its closest SA site (Figure 5D, Sup Figure 6C).

### Cryptic splicing event characteristics when no preferred SS available

The presence of multiple canonical SS within P15 seems to drive the high level of observed cryptic splicing outcomes whereas in P13 the cryptic splicing events are less frequent and more opportunistic. When looking at all the observed splice events within the S protein sequence, no obvious location patterns emerge – events can start and end across the entire span of the sequence (Figure 6A, Sup Figure 7A). While the majority of the splicing events relied on canonical SD (’GT’) and SA (’AG’) sequences or at least one of the two (yellow, Figure 6B), many events also displayed the use of non-canonical sequences (Figure 6C, Sup Figure 7B-C). These unusual splicing events are consistent with published data from endogenous genes (22). The length distribution of observed cryptic introns spans across the length of the transgene with a higher number of smaller events (Figure 6D, Sup Figure 7D-E). Splicing events with overall similar characteristics (Sup Figure 8) were also observed by Nanopore direct RNA sequencing in HEK293 cells transfected with ChAdOx1 nCov-19 vaccine vector (20) although reported at 4.2% of the full-length population. Similar splicing events were detected by cloning and sequencing RT-PCR transcripts following transfection of S protein sequences used in the CanSino (P18) and Sputnik V (P19) vaccines (Sup Figure 9A) even though CanSino S protein sequence design included removal of SS (2). Although the CanSino S protein sequence had the lowest average cryptic SS prediction score (4.3) it still contained two predicted ‘constitutive’ sites compared to 7-8 present in other codon-optimized S protein sequences (Sup Table 1).

**Figure 7.**
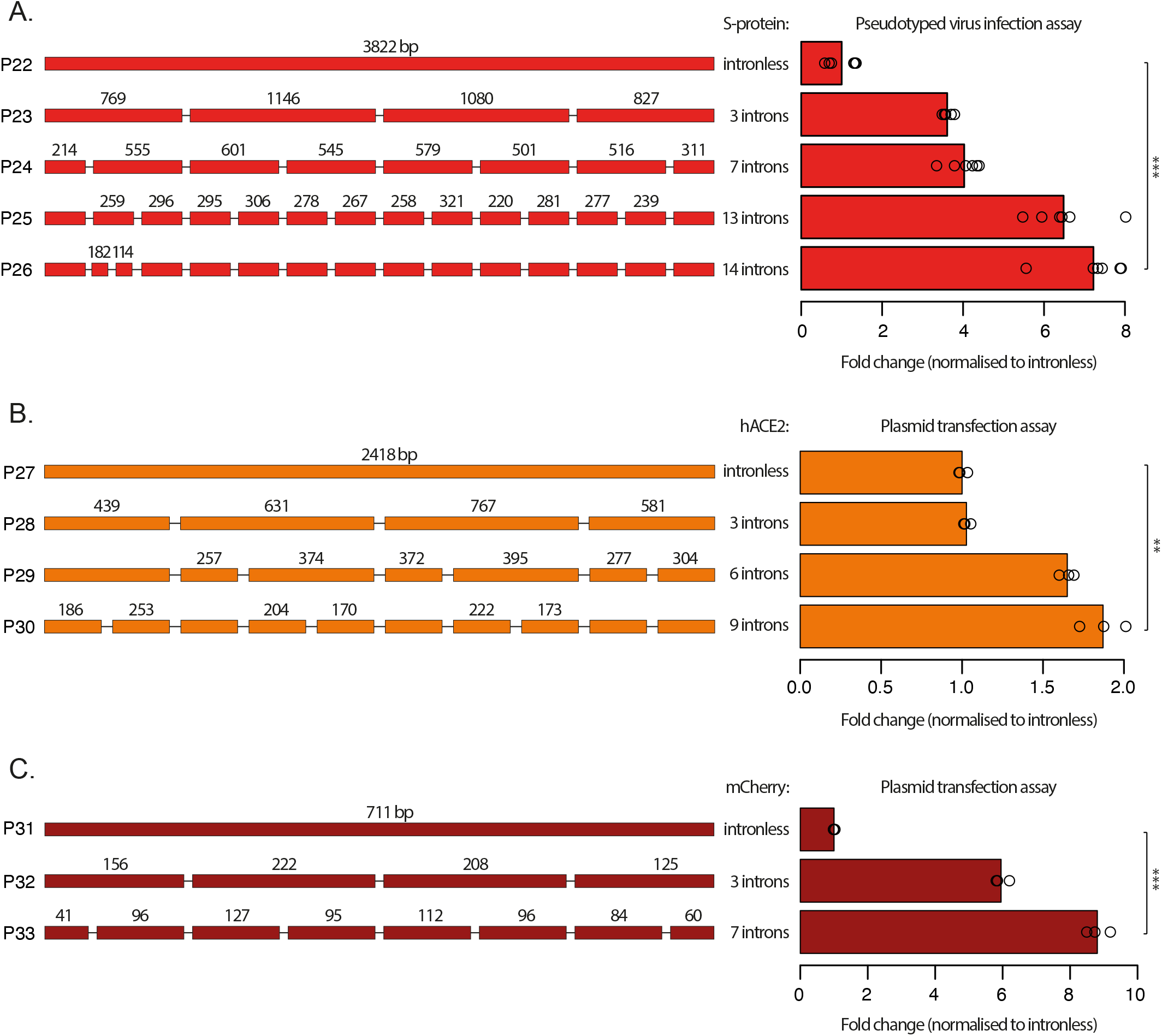
Intronization of S protein, hACE2 and mCherry transgenes improved protein expression. Four constructs with increasing number of introns (3–14) were introduced into S-protein CDS. Addition of more introns gradually improved protein expression and performance in pseudotyped virus infection assay **[A]**. The same trend was observed with 3 constructs containing increasing number of introns (3–9) introduced into hACE2 CDS **[B]** and 2 constructs (3,7) introduced into mCherry CDS **[C]**. **p<0.005, ***p<0.0005 (student t-test)

### Cryptic splicing is a general problem for transgenes

To assess splicing in other transgene settings, a codon-optimized viral sequence from the S protein of ChAdOx1 MERS vaccine candidate (P20) and a non-optimized human cDNA sequence for ACE2 protein (P21) were tested. Both of these constructs exhibited cryptic splicing (Sup Figure 9B-C). Similar cryptic splicing events were found in 8% of the published Nanopore RNA direct sequencing data from a cell line transfected with a construct designed to express a human cDNA encoding the ALKBH5 protein (23). The majority of the cryptic events stemmed from a few preferred canonical SS present within the transgene sequence (Sup Figure 10). All these data support the conclusion that observed cryptic splicing events are widespread and can be present in different transgenes regardless of the origin and optimization status of the CDS.

### Reconstruction of gene landscape by adding multiple introns improves transgene expression

Removal of canonical SS within transgene sequences can clearly alleviate repetitive cryptic splicing outcomes and improve transgene expression, exemplified in the performance difference between P13 and P15. However, this approach is insufficient in resolving the random cryptic splicing that continues despite the lack of optimal SS. This seemingly opportunistic splicing brought our attention to an obvious difference between transgenes and endogenous genes – the exon length.

While transgenes are typically designed as one long exon with an optional small 5’ non-coding exon, an average endogenous CDS in the human genome is broken up by more than 10 introns with an average length of coding exons at around 160 bp (24). SARS-CoV-2 S protein CDS is over 20 times longer than the exon length anticipated by the endogenous expression and splicing machinery. The introduction of small intronic sequences with conserved canonical SS at anticipated intervals was hypothesized as a potential solution for shifting random opportunistic splicing towards a controlled splicing outcome. To test this idea, several small endogenous human introns were introduced into various transgene CDS to achieve a more ‘natural’ gene landscape.

A number of criteria were considered for the selection of appropriate introns and their insertion sites within the transgenes to ensure correct recognition and splicing outcomes. To reduce possible confounding effects, all introns were initially chosen from a single human gene *Titin (TTN*) with the largest number of introns in human genome. All introns had to be small (80-92 bp) in order to have minimal impact on the overall length of the constructs. Introns were further prioritized to start with a good complementary sequence to the small nuclear RNA U1 (5′-ACUUACCUG-3′) responsible for the interactions with the donor site (25) and end with the canonical acceptor site ‘AG’. The chosen introns were inserted within the transgene cDNA sites with the sequence (C/A)AG || (G/N)(T/N)(T/N) or within sites where codon optimization allowed creation of such sequence without changes to the protein to further accommodate optimal binding of U1 and reflect the more commonly preferred SA sequence in human genome (26).

To assess the impact of intronization on top of standard codon optimization, the best expressing S protein sequence (P16) was used. An increasing number of introns were added and the 5’UTR intron as well as the 3’FLAG elements were removed (P22-P26, Figure 7A). Improvements in S protein expression were measured using a pseudotyped virus infection assay. This assay requires formation and function of an S trimer and is expected to more faithfully reflect levels of intact and functional S protein expression. The addition of introns and concomitant decrease of exon length, progressively improved S protein function as the number of introns increased resulting in a > 7-fold improvement for the best sequence (t-test: p<0.0005, Figure 7A). The maximum number of inserted introns (13 introns, exons ranging from 220-321 bp) was guided by the median (278 bp) and mean (321 bp) lengths of coding exons of *TTN* and were inserted in the 5’-3’ order of appearance within the *TTN* gene.

In order to assess whether the same principles apply in other transgenes, an increasing number of introns were next introduced to the hACE2 CDS, using some of the endogenous intron locations. While keeping the average exon size similar to *TTN*, two introns from other human genes (*RIF1* and *CCNG2*) were used here to demonstrate that intron origin is not relevant. Significant improvement in expression (t-test: p<0.005) was observed when 9 introns were introduced to the hACE2 sequence (P27-P30, Figure 7B).

To explore if further reduction to exon size can be beneficial, we introduced introns into the commonly used fluorophore mCherry. Despite mCherry CDS being only 711 bp long, intronization with 7 introns (exons ranging from 84 bp to 127 bp) increased average expression levels by approximately 9-fold (t-test: p<0.0005, P31-P33, Figure7C). These data demonstrate that intronization benefits expression of both endogenous as well as exogenous sequences. Implementation of this strategy may provide a general solution for managing cryptic splicing events and further improve protein expression from codon-optimized transgenes.

## Discussion

The coding elements of SARS-CoV-2 and other cytoplasmic viruses are optimized through Darwinian selection to serve as efficient templates for protein translation, replication and packaging. These sequences have not been subjected to selective pressures required for nuclear expression. Hence, expression of a SARS-CoV-2 gene CDS from a cDNA template in a nucleus will be inefficient. Potential cryptic SS, undesirable GC content, unintended transcription factor binding sites and other DNA features impacting sequence secondary structure or epigenetic landscape will all impact the performance of the sequence.

To achieve “efficient” expression of a CDS from a cytoplasmic RNA virus from a cDNA template, the rigorous iterative process of evolutionary selection may be substituted by a process of codon optimization. Through this process we adapt the sequence to conform to our understanding of the requirements of nuclear expression, capitalizing on the degenerate codon usage available for most amino acids. In principle the underlying DNA coding sequence can be adjusted by more than 65% without altering the underlying amino acid sequence. As codon usage *per se* is likely not the primary challenge for mammalian viral sequences which have evolved to use the same translational machinery, it really would be more pertinent to call this process sequence optimization. While codon optimization strategies do not directly target SS, GC content or other possible functional sequences, changing the sequence to the more frequently used codons may tackle these problems indirectly and inadvertently.

Analysis of the S protein wildtype sequence demonstrated that both cryptic SS and suboptimal GC content were critical obstacles for achieving full length RNA expression. While the wildtype S sequence (P1, P2) had strong preference of SS-usage, removal of these sites without really improving its GC content (P1: 37%, P12: 39%) was not sufficient for the sequence to be recognized as “exonic” by the expression and splicing machinery.

Cryptic splicing events still occurred in constructs where all canonical SS had been removed (P12) but they just became non-repetitive as a result of the use of random non-canonical SS. It is well-known that the exonic regions in humans and other mammals have a higher GC content (27) and inadvertently, the most used codon triplicates tend to end with either a ‘C’ or a ‘G’ where possible (Kazusa Codon Usage Database). Swapping most of the codons to the most common or second most common triplet significantly increased S protein sequence GC content (P13: 57%, P14: 55%, P16: 54%, Sup Table 2). Distributing codon usage equally between all options also improved GC content (P15: 46%, Sup Table 2). While increasing sequence GC content correlates with improved transcription as well as translation, high translation speed is not always considered optimal (28). In fact, many membrane proteins include less frequently used codons which is assumed to reduce the translation rate to facilitate correct folding (28). While the P15 optimization strategy may have benefitted S protein folding, the lack of attention to SS meant it was the least optimal codon-optimized S protein sequence we investigated.

Recent advances in long-read sequencing technologies such as Oxford Nanopore and PacBio have enabled us to capture splicing events better than ever before. The major advantage of the Nanopore cDNA direct sequencing approach used here was the lack of DNA amplification steps while still being able to enrich the sample with gene specific primers. The main limitation lies in the quantification of the observed events in comparison to the reads without events which can be influenced by size dependency of some of the sample preparation steps as the full-size S protein cDNA is over 4kb long and many of the observed splicing events can be shorter than 200 bp and depleted by many sample preparation methods. Reported here are distributions based on the full length read population alone within each sample, but care should be applied before comparing these frequencies to other datasets with different sample preparation methods, such as the ChAdOx1 Nanopore RNA direct data (Sup Figure 8).

Hundreds of randomly spaced splicing events characterized by the cDNA direct sequencing approach for the S protein and similar events seen in other proteins highlight the fact that even when the transgene’s CDS is apparently well designed and expression of full-length protein can be detected, the *status quo* design is just not optimal for RNA expression as transcript heterogeneity will inevitably impact both product levels (yield) and homogeneity. The impact of this will be dependent on the nature of the product, whether it is expressed *in vitro* or *in vivo*. *In vitro* expression offers opportunities to improve homogeneity by application of purification methods. Two thirds of cryptically spliced RNA molecules would be out-of-frame and would not just impact protein yield but any translation *in vivo* would generate novel peptides with potential immunogenicity. In-frame products are potentially of greater impact. In an *in vitro* expression system, they may co-purify with the full-length product requiring more complex purification processes to achieve a homogenous product. Proteins like the trimeric S-protein are particularly vulnerable to disruption by in frame fragments as these can be incorporated into, and have dominant negative impact on the complex as exemplified by human germ line mutations in trimeric collagens causing diseases like Osteogenesis Imperfecta (29).

While the benefit of having a 5’UTR intron was identified decades ago (8), frequent introns spaced throughout the gene were not considered as practical or necessary. The genes for many human therapeutic proteins cover large genomic expanses and contain numerous large introns. Such loci are significantly larger than the cloning capacity of vectors used for expression purposes and therefore these have been most conveniently handled as cDNAs. While in many cases these do express at some level, the benefits that introns bring has been under appreciated compared to the other factors that impact expression such as copy number, promoter strength and integration site. Furthermore, the epigenetic landscape of exons and introns was not well understood, even though intron/exon boundaries are highly conserved in orthologous and paralogues across hundreds of millions of years of evolution.

Regenerating a more ‘natural’ landscape in transgenes had an immediate impact on protein expression and function. Introns improved expression in the context of sequences that naturally do not harbor introns to begin with (S protein), as well as when inserted into non-endogenous locations (mCherry) or endogenous intron locations with no change to the native exonic sequences (hACE2). The latter highlights how intronization approach can serve as an alternative way of increasing recombinant protein expression without the need to change the natural codon selection in products where codon optimization has been shown to impact protein structure and function.

Intron-exon landscape reengineering not only guides the splicing events towards specific splicing outcomes, but may also play a role in increasing recognition, accessibility and/or recruitment of expression machinery – an area that will need further investigation in the future. The multi-fold expression improvement and anticipated reduction in product heterogeneity has wide implications within the medical field alone from recombinant protein production to vaccines and *in vivo* therapies.

## Methods

### DNA plasmids

Table 1 gives an overview of all the DNA plasmids expressing SARS-CoV-2/MERS Spike and hACE2 protein used in this paper. Full DNA sequences of these plasmids are found at a Zenodo provided doi: 10.5281/zenodo.5470001. “Wuhan” in plasmid names refers to the S protein DNA sequence from the Wuhan-Hu-1 isolate (Genbank: MN908947.3) while “18F” refers to the removal of the last 18 amino acids of the S protein C terminus (ER retention sequence) and the addition of a FLAG tag.

Plasmid P2, a commercially available S protein expression vector based on the Wuhan-Hu-1 isolate, was purchased from InvivoGen (pUNO1-SARS2-S(D614)). All other plasmids were assembled by Gibson cloning of relevant PCR products and/or custom designed gene blocks (gBlocks Gene Fragments, Integrated DNA Technologies) according to manufacturer’s protocol (Gibson Assembly Master Mix, NEB). The template plasmids for the codon-optimized SARS-CoV-2 S protein sequences were generously shared with us by Professor Robin Shattock from the Imperial College London (S-Imperial), Dr Nigel Temperton from the University of Kent (S-Kent), by Professor Nevan Krogan from the University of California San Francisco (S-UCSC) and by Dr Emma Bentley from the National Institute for Biological Standards and Control (S-NIBSC), respectively. The DNA sequences for the codon-optimized SARS CoV-2 S protein used in CanSino and Sputnik V vaccines were obtained from relevant patents (CN110974950B, RU2731356C1). The DNA sequence for the codon-optimized MERS-CoV S protein used in the Oxford ChAdOx1 vaccine candidate was obtained from the relevant patent (WO2018215766A1). mCherry CDS refers to the “synthetic construct monomeric red fluorescent protein gene” (Genbank: AY678264.1), with the stop codon changed from ‘TAA’ to ‘TGA’ while human ACE2 CDS refers to the “Homo sapiens angiotensin converting enzyme 2, mRNA transcript variant 2” (Genbank: NM_021804.3).

### *In vitro* transcription of S protein mRNA

Templates for *in vitro* transcription were generated by cloning P1 and P13 between NcoI and NotI sites of pTNT-B18R-6His (addgene plasmid 58979, a kind gift from Steven Dowdy (30)). Plasmids were linearised through BamHI digestion and purified with Agencourt AMPure XP beads. 2 ug were used as template for *in vitro* transcription with mMESSAGE mMACHINE T7 ULTRA Transcription Kit (Ambion), according to the manufacturer’s instructions. Following polyadenylation step, mRNA was purified using MegaClear kit (Ambion) and eluted in 100 μL of RNase-free water. Produced mRNA was then concentrated by precipitation with 5M ammonium acetate and absolute ethanol. mRNA quality was assessed by denaturing agarose gel electrophoresis and nanodrop quantification.

### Cell lines used in this study

293FT cells were obtained from Dr. Kosuke Yusa’s Lab. 293FT.Cas9 cell lines were generated through lentiviral integration of an EF1a-Cas9-T2A-BlastR construct at low MOI to achieve single-copy integration. To generate cell lines permissive to Spike-Pseudotyped lentiviral infection, 293FT.Cas9 cells were engineered to stably express SARS-CoV-2 receptors ACE2 and TMPRSS2. Stable expression was achieved by PiggyBac transposition of a EF1a-ACE2-T2A-TMPRSS2 constructs, followed by single-cell cloning. This resulted in 293FT.Cas9.ACE2/TMPRSS2 clonal cell lines. Clones C10 and D10 were used in this work. 293T cells were obtained from Dr. Ravindra Gupta Lab (University of Cambridge) and used for Spike-Pseudotyped lentivirus production. All cell lines have tested negative for mycoplasma contamination.

### Cell culture conditions

Unless stated otherwise, all cell lines were maintained at 5% CO2 and 37°C. All cell lines were routinely cultured in M10 media (DMEM, 10% FBS and 2 mM L-glutamine). Cas9 expressing cell lines were maintained in M10 supplemented with 10 μg/mL Blasticidin.

### Cell transfections

Cell transfection was carried out using Lipofectamine LTX Reagent (Invitrogen) according to the manufacturer’s instructions. For 6-well format transfections, Lipofectamine::DNA complexes were formulated using 750 ng DNA, 5 μL Plus reagent and 10 μL Lipofectamine LTX. These were then used to transfect 1.5 million cells per reaction. For analysis of transgene expression, cells were harvested with trypsin generally 48h post-transfection. Samples were kept as frozen cell pellet for cDNA analysis, as whole-cell extract in RIPA buffer supplemented with protease inhibitors for Western Blot analysis, or used directly for flow cytometry assays. For *in vitro* transcribed mRNA, 293T cells were transfected for functional testing using Lipofectamine messengerMAX (Invitrogen) according to the manufacturer’s instructions. Briefly, for 24-well format, 625 ng of S protein mRNA were incubated with 0.75 ul of Lipofectamine messengerMAX and used to transfect 0.3 million cells per reaction.

### Flow cytometry assays

Cells were harvested 48h post-transfection, using trypsin dissociation. Upon harvesting, cells were washed twice with staining buffer (see Table 2). They were then incubated with the appropriate dilution of primary antibody (in staining buffer) for 30 min at the indicated temperature. Cells were washed twice and incubated with secondary antibody (1:500) for 30 min on ice (for non-conjugated primary antibodies). Following another set of two washes, cells were analysed by flow cytometry using Cytoflex (BD Biosciences). Data analysis was performed using FlowJo software (BD Biosciences). S protein expression data is plotted as % of positively stained cells (Figure 1, 4, Sup Fig 1), while ACE2 and mCherry data is visualized as intensity median value normalized to intronless constructs to highlight the shift in population intensity (Figure 7).

### Pseudotyped Lentivirus productions

Pseudotyped Lentivirus was produced by transfection of 293T cells using lipofectamine LTX according to the manufacturer’s instructions. Intronized S protein constructs were tested using three independent virus productions. Briefly, 1 million 293T cells were seeded into gelatinized 6-well plates one day ahead of transfection. For transfection, 1 μg of lentiviral transfer vector (pCSGW-GFP), were mixed with 0.72 μg of gag-pol expressing plasmid p8.9 and 68.33 fmol of S protein expressing construct in 500 μL of optiMEM media followed by the addition of 2 μL of PLUS reagent and incubation for 5 minutes at room temperature. 6 μL of Lipofectamine LTX reagent were then added to the mix and incubated for 10 minutes. Medium was aspirated from the plates and the Lipofectamine:DNA complexes were added dropwise and topped-up with 1.5 mL of M10. Production was carried out at 5% CO_2_ and 32°C. Medium was changed the following morning to 2.5 mL of fresh M10 and the supernatant was harvested 56 hours later. Virus-containing supernatant was spun down at 500g for 5 minutes to remove cell debris and used directly to infect permissive cell lines or aliquoted and frozen at −80°C.

### Permissive cell line transduction

Transductions were carried-out in 96-well plates, in duplicates for each independent virus sample. For pseudotyped lentivirus titrations, a dilution series was prepared ranging from 100% virus-containing supernatant to 1:500 dilution in a total volume of 200 μL M10 medium. 293FT.Cas9.ACE2/TMPRSS2 clonal cell lines were harvested by trypsinization and resuspend at a density of 70.000 cells per 30 μL. They were then seeded, 30 μL per well, mixed and incubated at 37°C. Viral infection efficiency was measured 48-72h later, assessed by the percentage of GFP positive cells on flow cytometry. Data was analysed using FlowJo software (BD Biosciences) and displayed as % cells infected at 1:500 dilution of pseudotyped virus, normalized to the intronless construct infection rates (Figure 7).

### Western Blot

Preprocessing of samples included sonification, separation from cell debris by centrifugation, assessment of protein concentration using Pierce™ BCA Protein Assay Kit (Thermo cat#23225) and addition of 4x Laemmli Sample buffer (Bio-Rad). 40ng of each sample was loaded onto Mini-PROTEAN GTX Precast Protein Gel (Bio-Rad, #1610732) together with Precision Plus Protein Dual Color Standard Ladder (Bio-Rad, #1610374). Trans-Blot Turbo Transfer System (Bio-Rad) was used to dry transfer the material onto a PVDF membrane (Trans-Blot Turbo Mini 0.2 ul PVDF Transfer Packs, Bio-Rad, #1704156). After blocking with 5% skim milk in TBS-T, incubations with primary and secondary antibodies were carried out as detailed in Table 3. Staining was developed using 20X LumiGLO® Reagent and 20X Peroxide reagents according to manufacturer’s recommendations (Cell Signaling Technology, #7003).

**Table 3.** Western blot antibodies.

| **Antibody** | **Source** | **Conditions** |
| --- | --- | --- |
| anti-S, mouse IgG1 | Genetex #GTX632604 | 1:2000 dilution, 4°C, overnight |
| anti-FLAG, mouse IgG1 | Sigma-Aldrich #F1804 | 1:4000 dilution, 4°C, overnight |
| anti-GAPDH, mouse IgG1 | Abcam #ab8245 | 1:2000 dilution, 4°C, overnight |
| Goat Anti-mouse IgG conjugated with HRP | Abcam, #ab205719 | 1:10000 dilution, RT, 1h |

### cDNA analysis

RNA was extracted from the frozen cell pellets using RNeasy Mini Kit (Qiagen) and treated with ezDNase (ThermoFisher) before applying oligo(dT) guided 1^st^ strand cDNA synthesis using SuperScript IV reverse transcriptase (ThermoFisher), all according to manufactures’ recommendations. RT-PCR was carried out using GoTaq Green Master Mix (Promega), following recommended protocol. PCR primers used to capture the entire length of investigated transgenes are enlisted in Table 4.

**Table 4.**
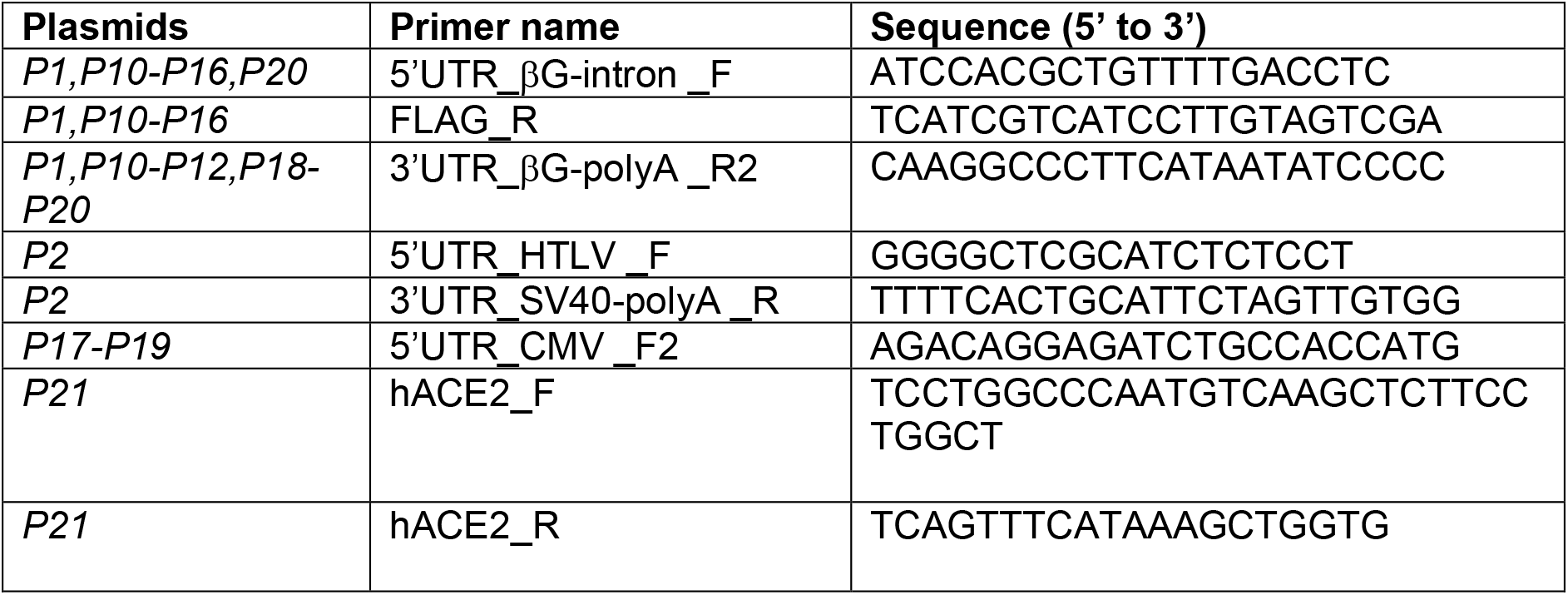
RT-PCR primers

These PCR products were both visualized on an agarose gel as well as TA-cloned using ‘TA Cloning Kit with pCR2.1 vector and OneShot TOP10 Chemically Competent E.coli’ (ThermoFisher) according to kit instructions. After overnight growth on LB plates containing 100 μg/ml ampicillin at 37°C, single colonies were picked into 20 μl of PBS and the respective vector insert was PCR amplified with M13F (GTAAAACGACGGCCAGT) and M13R (CAGGAAACAGCTATGAC) primers, using GoTaq Green Master Mix. These PCR products were purified using AmPure XP magnetic beads (Beckman Coulter) following manufacture’s recommendations and submitted to Sanger Sequencing (supplied by Source BioScience Inc) using the above M13F/M13R primers. On average, 24 clones per construct were assessed by PCR and 8 clones further selected for Sanger sequencing. All reads were mapped back to the original construct DNA sequence using SnapGene software to assess individual mRNA splicing events.

### Nanopore sequencing sample prep

The sequencing samples were prepared according to the Direct cDNA Sequencing protocol (Nanopore, SQK-DCS109) with a few modifications to increase the abundance of transcript of interest. RNA was extracted from the frozen cell pellets using RNeasy Mini Kit (Qiagen) and treated with ezDNase (ThermoFisher). Three independent 1^st^ strand cDNA synthesis reactions using SuperScript IV reverse transcriptase (ThermoFisher) were carried out using a transcript specific reverse primer (either FLAG_R or 3’UTR_βG-polyA _R2, Table 4). RNA was degraded using RNase Cocktail Enzyme Mix (ThermoFisher #AM2286). The three samples were next bead purified (AMPure XP beads) using three different volume ratios (1X-2X-4X) to ensure capturing the entire size range of splicing products. 2nd cDNA strand was again synthesized using a transcript specific primer (either 5’UTR_βG-intron _F or 5’UTR_CMV _F2, Table 4) and Q5 2x master mix (NEB, #M0494). After another round of bead purification, samples were pooled and submitted to Edinburgh Genomics, where they were further processed, barcoded and run on PromethION platform.

### Nanopore sequencing data analysis

Reads from all samples were aligned to the relevant construct templates using minimap2. Parameters for the different minimap2 alignments can be found here: https://github.com/jacobhepkema/RNA_deletions. The aligned reads were next investigated for splicing events using an in-house script available at the above repository. Reads for the ALKBH5 experiment were downloaded from SRA accession SRR9646144, and reads for the ChAdOx1 experiment were downloaded from SRA accession SRR13320597. The template used to align the ALKBH5 reads to was constructed manually, by performing *in silico* Gateway cloning, inserting the ALKBH5 CDS and mRuby3 CDS into the pLIX_403 vector (Addgene plasmid #41395). The linker region between ALKBH5 and mRuby3 was constructed manually by finding the consensus at each position for reads mapping to the end of the ALKBH5 CDS or the start of the mRuby3 CDS, with the requirement that the ALKBH5 CDS and mRuby3 CDS were in-frame. Overview of all the aligned reads, splicing events, and span is available in Sup Table 3. All data visualization was created using R (basics).

## Acknowledgements

This work was funded by UK Research and Innovation (BB/V011316/1). AB is further supported by a Wellcome Trust research grant (210525/Z/18/Z) and KT is the recipient of the Sir Henry Wellcome postdoctoral fellowship (Wellcome grant number: 210910/Z/18/Z). Nanopore cDNA direct library preparation and sequencing was carried out by Edinburgh Genomics.

## Author Contributions

Conceived and designed experiments: KT, AB. Construction designs, cloning, cDNA preparation and analysis: KT. Tissue culture, mRNA synthesis, flow cytometry and pseudotyped assays: LA. Western blots: YP. Nanopore sequence sample prep: KT, DG. Computational analysis: KT, JH, AM. Wrote the first manuscript: KT, AB with input from all authors.

---

**Supplementary Figure 1.**
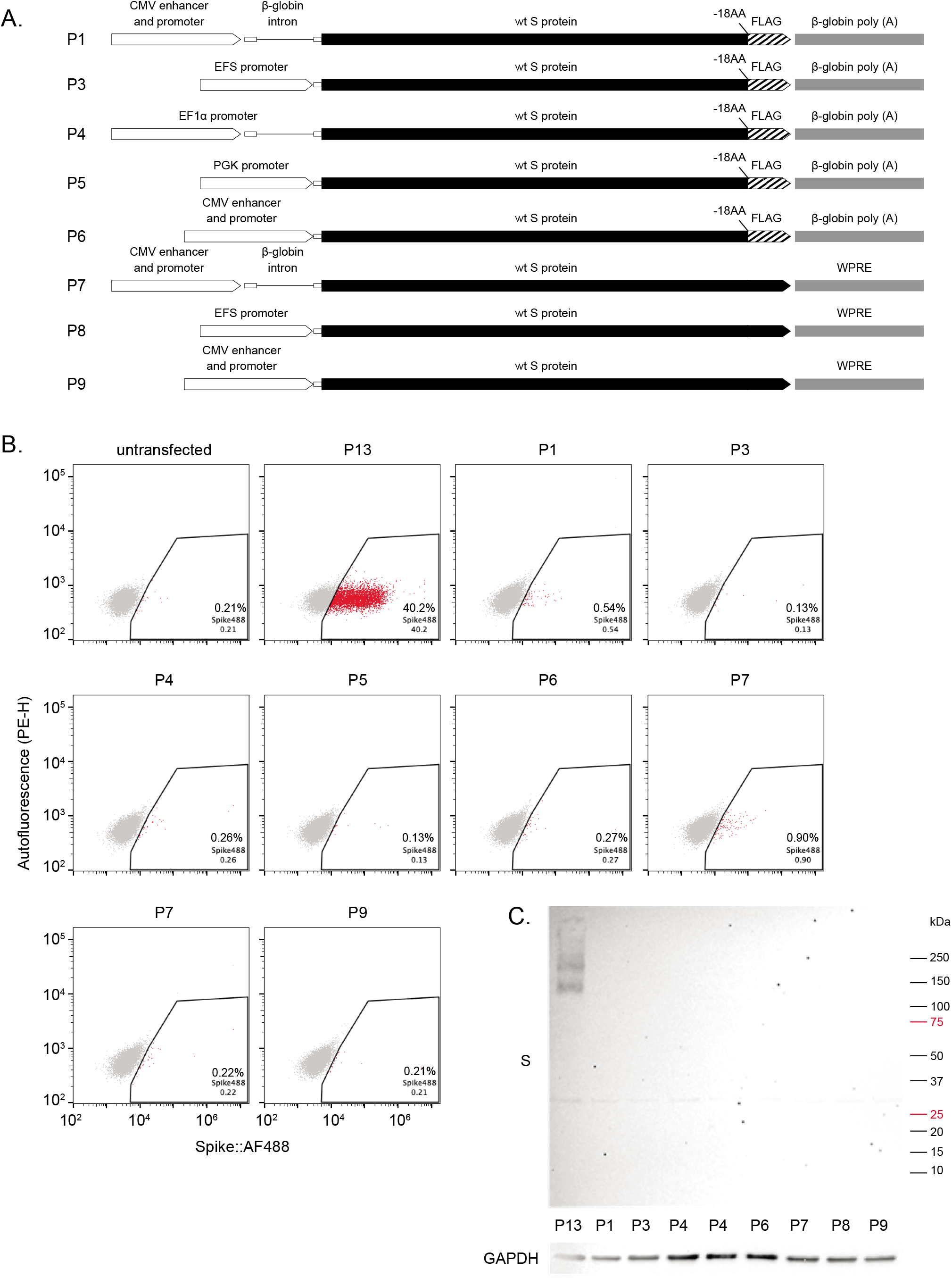
Swapping up- and downstream expression elements does not markedly change expression. Seven additional variations of the wt S construct were created by combining different up- and downstream elements, removing the FLAG tag and adding back the last 18AA to the S protein sequence **[A]**. None of these variations changed the expression level of S protein, measured both by flow cytometry **[B]** as well as Western Blot **[C]**.

**Supplementary Figure 2.**
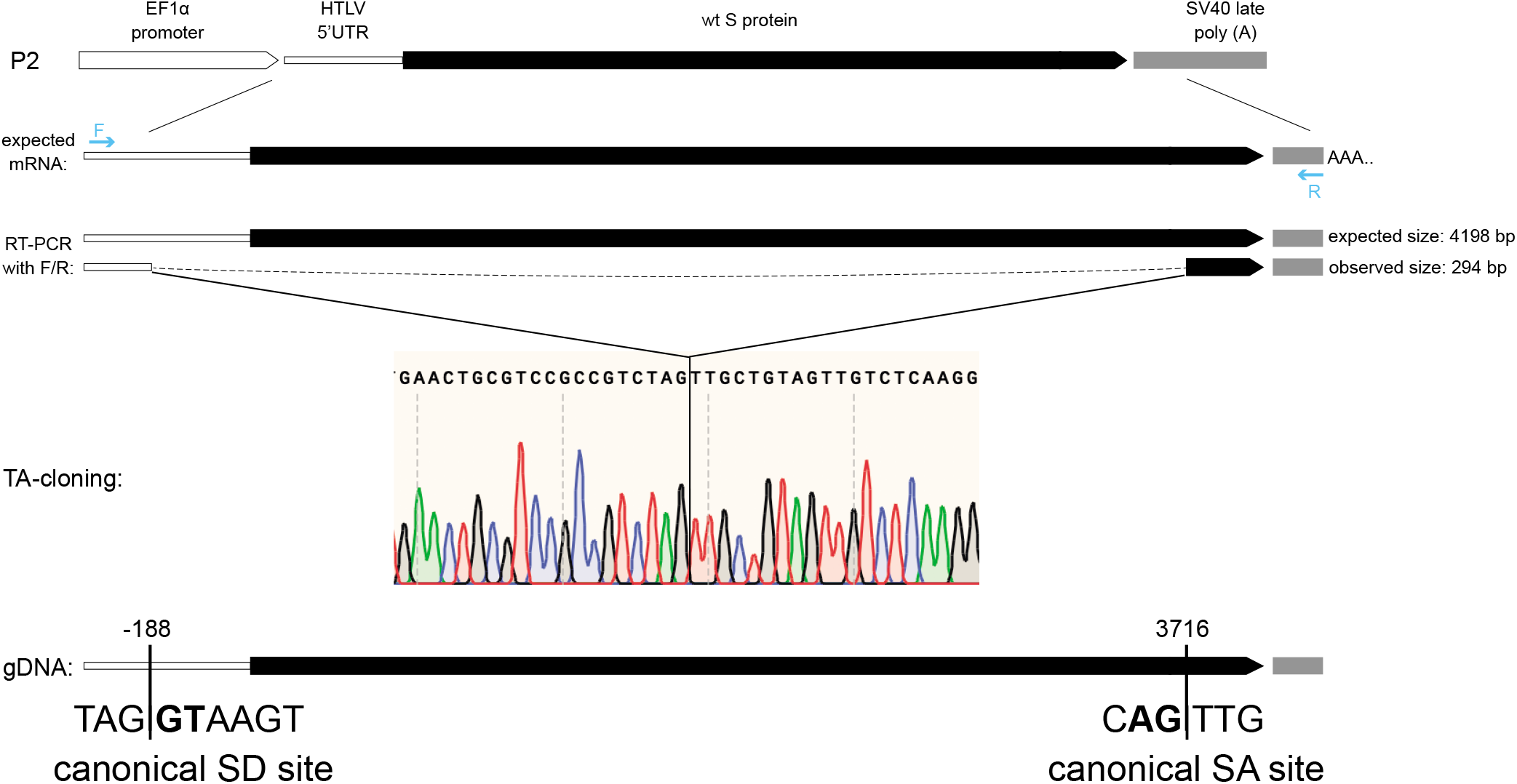
Commercially available wt S protein construct P2 also leads to cryptic splicing events. RT-PCR with primers positioned from 5’UTR to 3’UTR of the P2 construct (F,R) followed by TA-cloning and Sanger sequencing identified cryptically spliced mRNA product (294bp) that uses a canonical SD site within the HTLV 5’UTR (also used by HTLV within native setting) and a SA site available within the wt S protein sequence at position 3716 (same SA site as for P1).

**Supplementary Figure 3.**
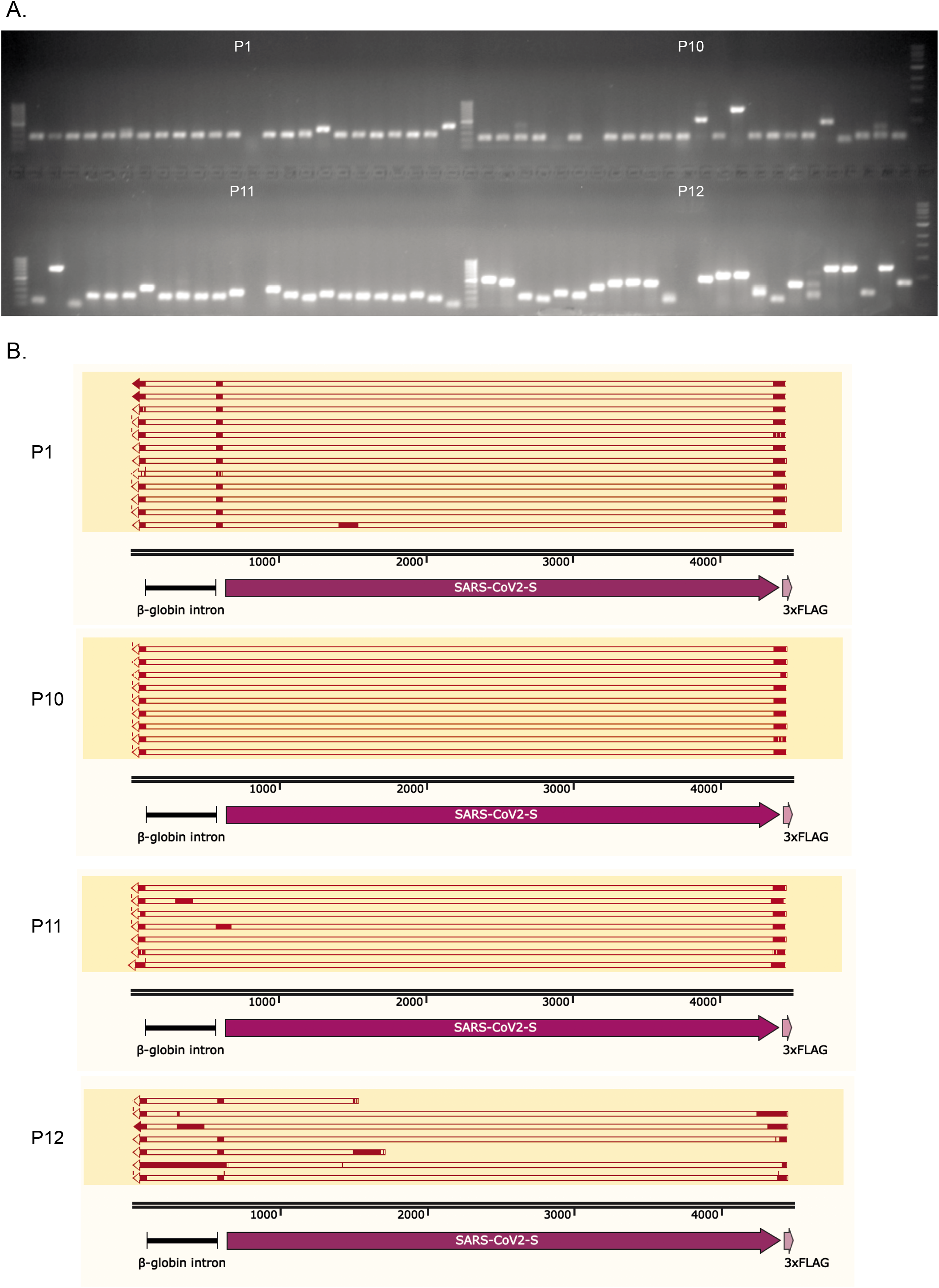
Removal of splice sites diversifies cryptic splicing events. A set of 24 clones was picked for each of the 4 constructs after TA-cloning the RT-PCR products with F/R primers (P1, P10-P12) **[A]**. Some of the clones were further Sanger sequenced and mapped against the relevant construct template **[B]**.

**Supplementary Figure 4.**
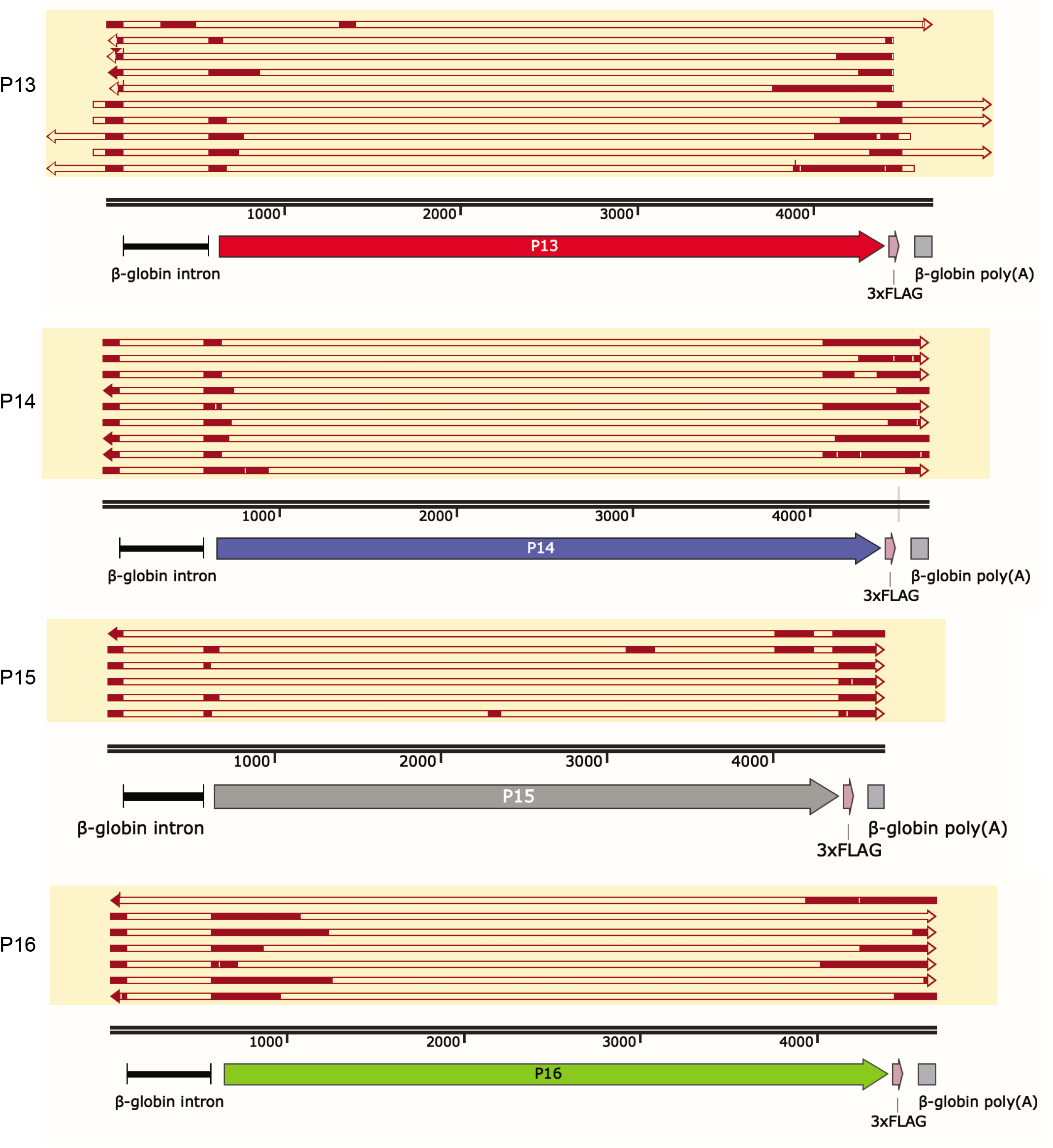
Examples of alternatively spliced products from c-o constructs. All c-o constructs expressed alternatively spliced products. In some constructs, such as P13 and P16, all splicing events captured by TA-cloning used different and not necessarily canonical SS while in P14 and P15 most alternatively spliced products used the same set of canonical SS.

**Supplementary Figure 5.**
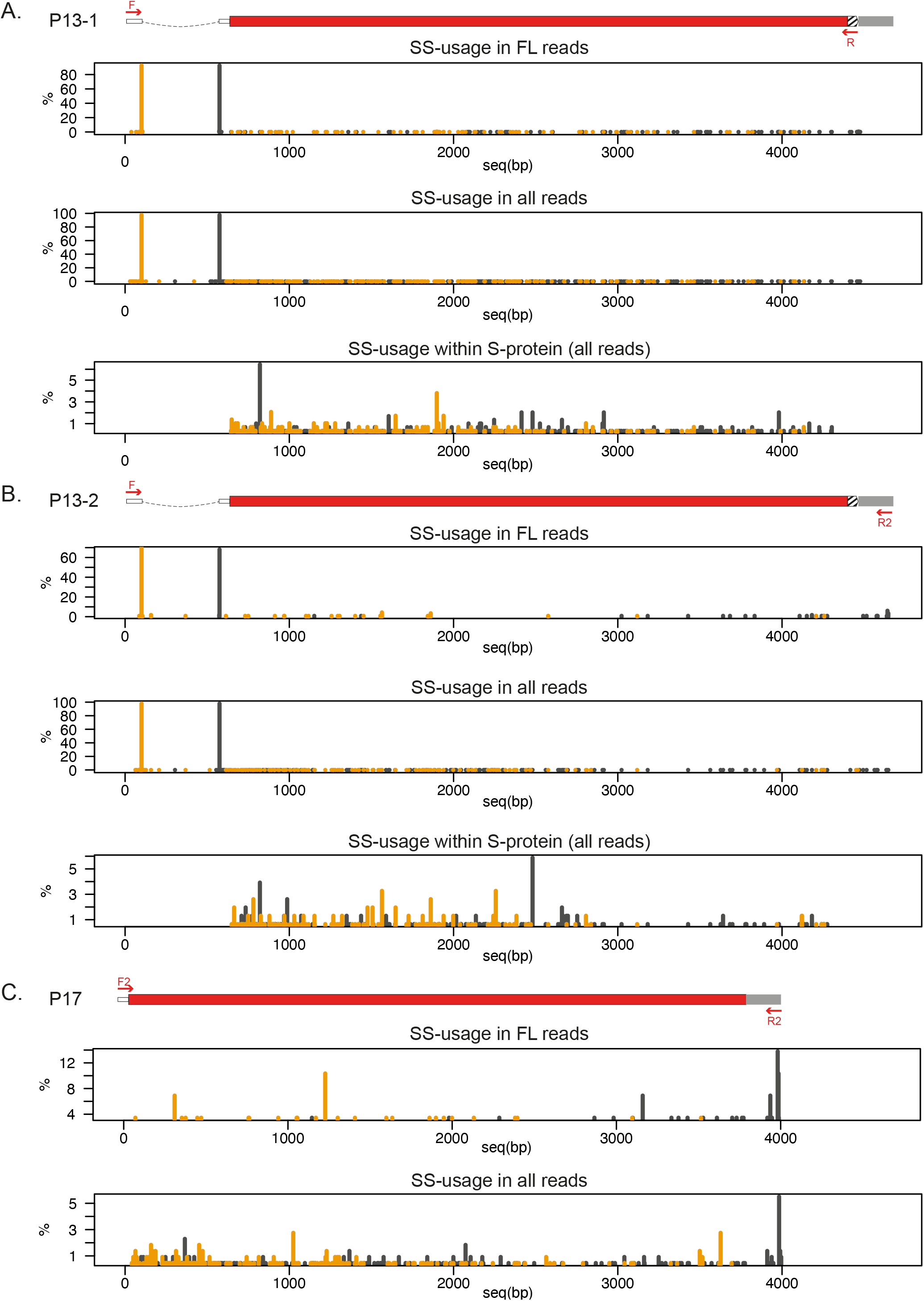
SS-usage characteristics for P13 similar across all datasets. The lack of strong preference in SS-usage observed in FL reads is retained when analyzing all the mapped reads in P13-1 **[A]** and P13-2 **[B]** samples, highlighted by displaying SS within only the S-protein CDS. Same SS-usage pattern remains in P17, a construct without either the 5’UTR intron or the −18AA,FLAG tag.

**Supplementary Figure 6.**
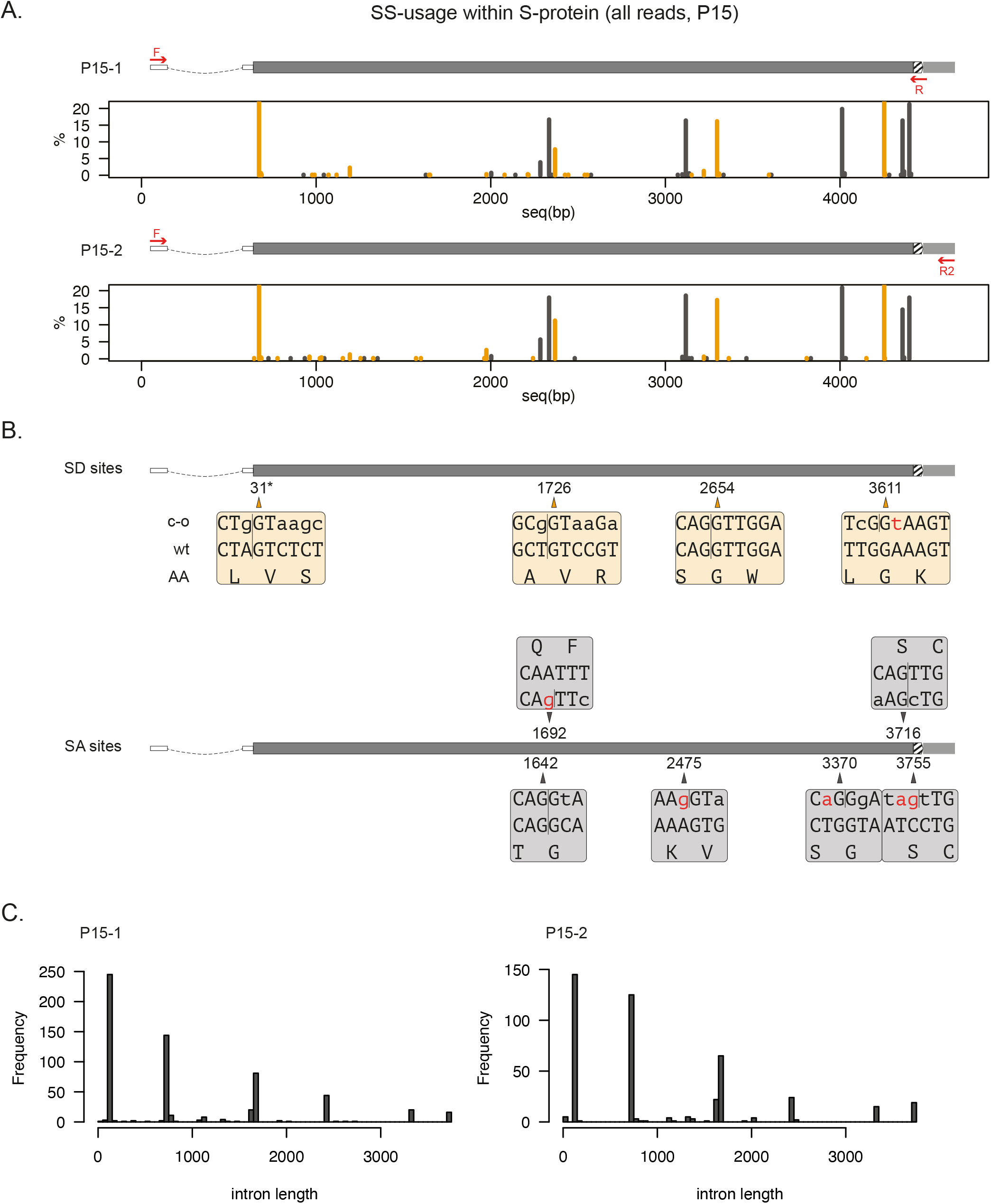
SS-usage characteristics for P15 similar across datasets. All the preferred SS observed in P15 FL reads (Figure 5) remain prevalent when looking at SS-usage within all reads (SS within S-protein displayed here) **[A]**. Some of these sites existed in the original wt S-protein sequence, while others were introduced during c-o. Lower case indicates nucleotide changes in c-o S, red highlights changes that resulted in novel cryptic SS **[B]**. Distribution across intron lengths for P15-1 and P15-2 samples **[C]**. (*) SS positions relative to translation start site ATG.

**Supplementary Figure 7.**
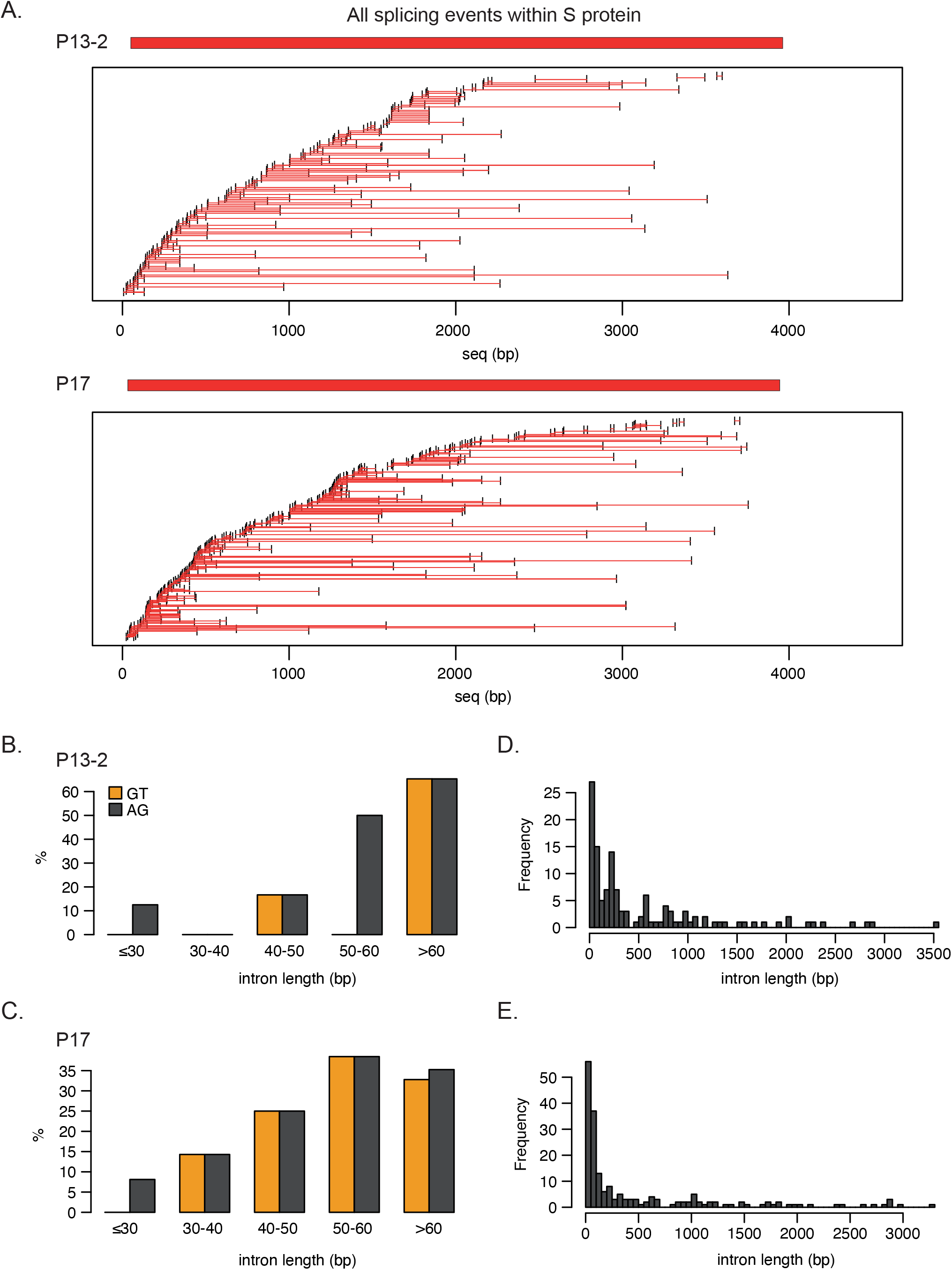
All P13-2/P17 splicing events within S-protein sequence similar to P13-1. P13-2/P17 splicing events are also mostly unique and spread across the S-protein sequence **[A]**. The canonical SS-usage increases for introns over 50bp both for P13-2 **[B]** and P17 **[C]** and the distribution of intron length for P13-2 **[D]** as well as P17 **[E]** remains consistent with P13-1.

**Supplementary Figure 8.**
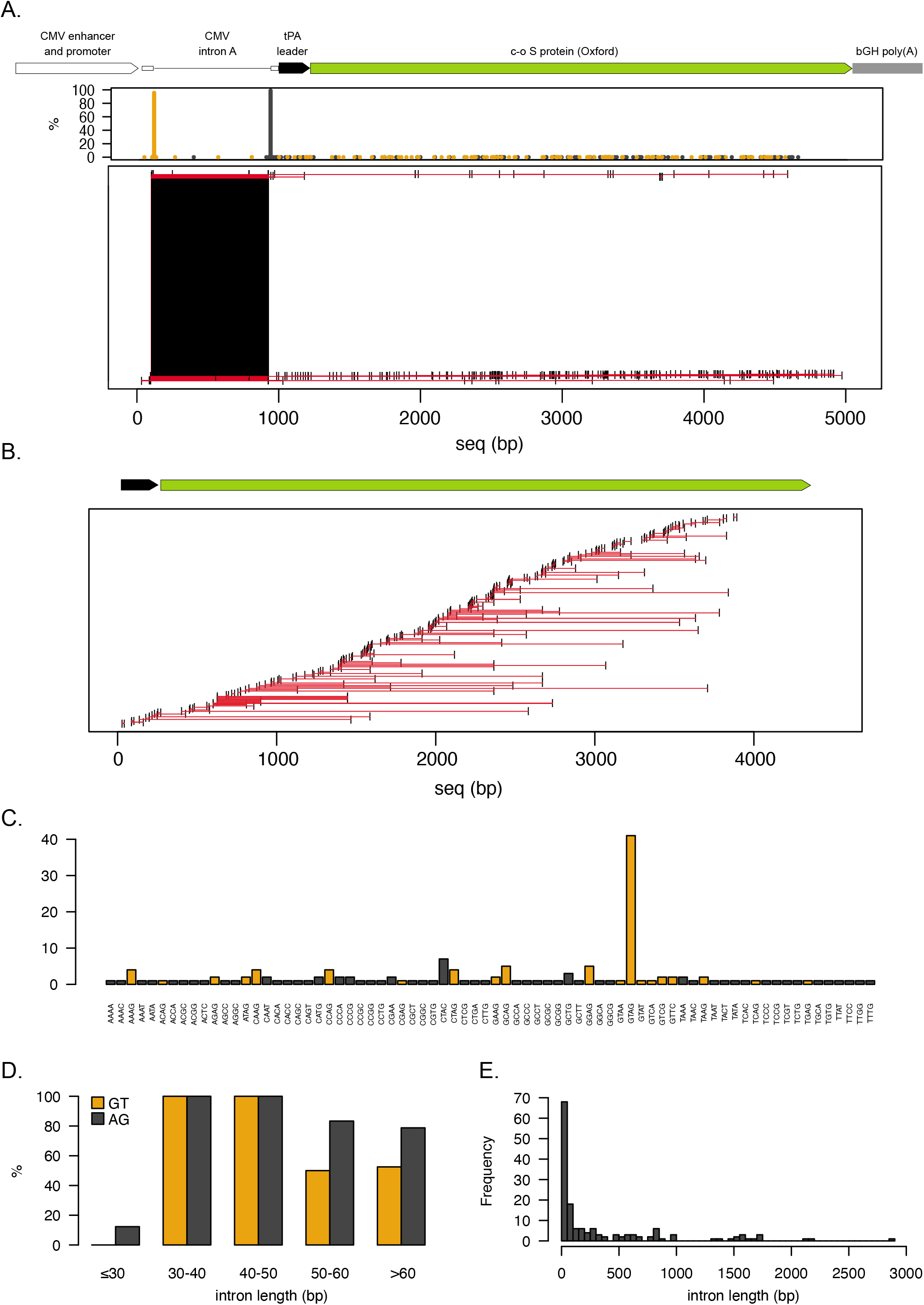
Cryptic splicing events in ChAdOx1 nCov-19 vaccine S protein. Overview of all identified splicing events **[A]** and all events within the S protein CDS **[B]**. SS-usage of all events within S protein **[C,D]** and the length distribution of the observed introns **[E]**.

**Supplementary Figure 9.**
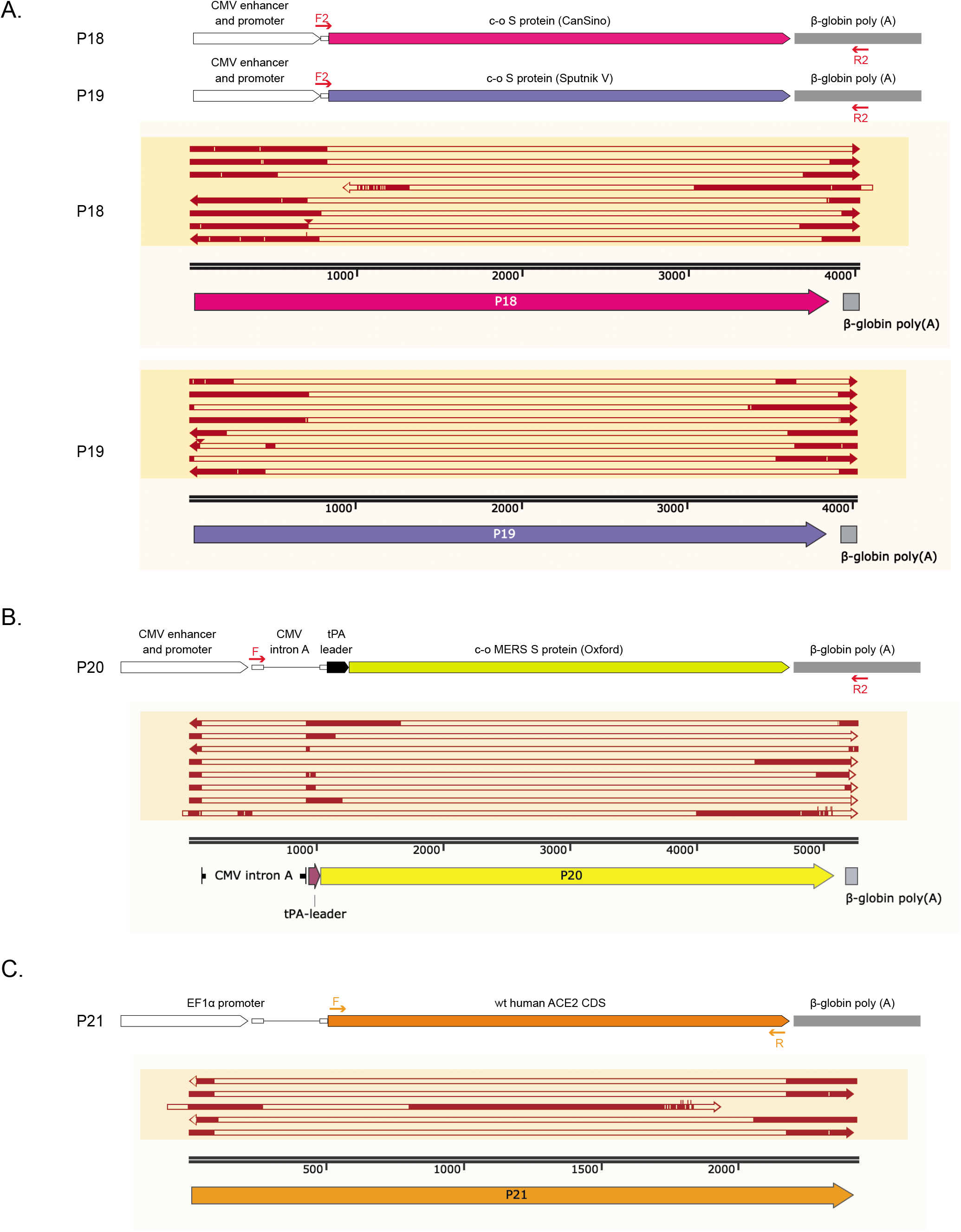
Cryptic splicing is a generic transgene problem. Similar splicing events to c-o S proteins were observed in currently available vaccine sequences CanSino and Sputnik V **[A]**. These events are not unique to SARS-CoV2 S-protein but also found in MERS vaccine sequence **[B]** as well as in human ACE2 CDS when expressed as a transgene **[C]**.

**Supplementary Figure 10.**
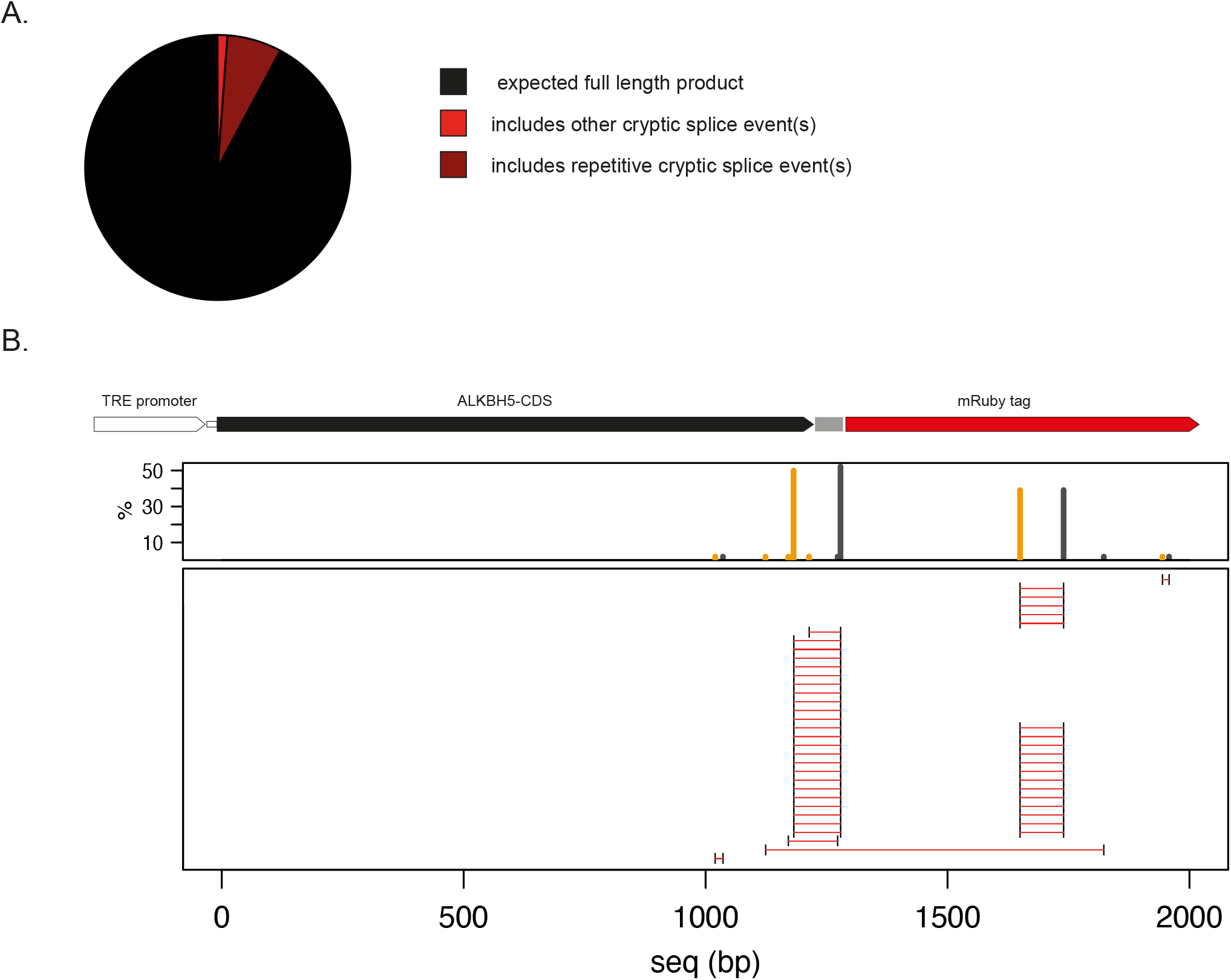
Cryptic splicing in *ALKBH5* transgene in HMEC cells. Approximately 8% of the FL reads aligned to ALKHB5 transgene contained a cryptic splice event **[A]**, with a preference of two predominant events **[B]**.

